# Toxicity of extracellular cGAMP and its analogs to T cells is due to SLC7A1-mediated import

**DOI:** 10.1101/2024.09.21.614248

**Authors:** Valentino Sudaryo, Dayanne R. Carvalho, J. Michelle Lee, Jacqueline A. Carozza, Xujun Cao, Anthony F. Cordova, Lingyin Li

## Abstract

STING agonists are promising innate immune therapies and can synergize with adaptive immune checkpoint blockade therapies for cancer treatment, but their effectiveness is limited by the toxicity to activated T cells. An important class of STING agonists are analogs of the endogenous STING agonist, cGAMP, and while transporters for these small molecules are known in some cell types, how they enter and kill T cells remains unknown. Here, we identify the cationic amino acid transporter SLC7A1 as the dominant transporter of cGAMP and its analogs in activated primary mouse and human T cells. T cells upregulate this transporter upon activation and rapid proliferation to meet their high metabolic demand, but this comes at the cost of enabling increased transport and toxicity of cGAMP. To circumvent the essentiality of SLC7A1 to proliferating T cells, we found that the residues responsible for cGAMP transport are separate from the arginine binding pocket allowing us to perturb cGAMP transport and STING-activation mediated killing without impacting arginine transport. These results suggest that SLC7A1 is a potential target for alleviating T cell toxicity associated with cGAMP and its analogs.

## Introduction

A hallmark of cancer is the presence of double-stranded DNA in the cytosol, resulting from chromosomal instability, damaged mitochondria, extrachromosomal DNA replication or treatment-induced chromatin bridges^1–4^. The innate immune pattern recognition receptor cGAS, upon recognizing cytosolic dsDNA, is activated to produce the second messenger cGAMP. cGAMP is a potent activator of its ER transmembrane receptor STING (Stimulator of Interferon Genes), which scaffolds the recruitment and activation of kinase TBK1 and transcription factor IRF3, which ultimately stimulates type-I interferon gene expression^5^. Cancer cells can avoid cGAMP-induced STING activation and interferon production by exporting cGAMP using transporters such as ABCC1^6,7^. Once released into the tumor microenvironment, cGAMP can enter bystander cells including tumor endothelial cells^8^, tumor-infiltrating macrophages and dendritic cells^6,9^, and tumor associated fibroblasts^10^ through cell type specific transporters. cGAMP-mediated activation of STING and type-I interferons in these cell types promotes immune infiltration into the tumor microenvironment^11,12^, producing an intercellular signaling cascade that can turn immunologically “cold” tumors “hot”.

cGAMP mimetics and traditional small molecule STING agonists were developed to improve the efficacy of checkpoint blockade therapies – which target the adaptive immune arm and therefore often fail to produce strong responses in immunologically “cold” tumors^13–16^. Injection of STING agonists into mouse tumors leads to remarkable shrinkage of these tumors and abscopal effects on distal tumors in a T cell-dependent manner^14,17^. Disappointingly, some studies have shown a loss of efficacy at higher dosages of these agonists, which has been attributed to toxic effects of STING activation in T cells recruited to the tumor microenvironment ^18–25^. This impact on T cells narrows the therapeutic window of and presents challenges for the further development and use of STING agonists. It may also underly limitations to the curative effect of radiation therapy due to endogenous cGAMP released into the tumor microenvironment.

Elucidating the cell type-specific cGAMP transporter(s) acting in human T cells could provide a means to selectively diminish the negative effects of cGAMP and other STING agonists on T cell survival and function, while maintaining their anti-cancer effects on other cell types in the tumor microenvironment. The LRRC8A:C heteromeric channel was reported to induce mouse T cell death by acting as a cGAMP channel to facilitate STING-P53 activation^20^. However, it is not known whether the LRRC8A:C channel transports cGAMP in human T cells, or whether it is in fact the dominant or only cGAMP transporter in mouse T cells.

In this study, we show that, the cationic amino acid transporter SLC7A1 is the dominant T cell-specific cGAMP transporter in activated mouse and human T cells. SLC7A1 is highly upregulated in activated T cells to transport the essential amino acid, arginine, but also transports endogenous cGAMP secreted by cancer cells and synthetic cGAMP analogs. Therefore, this upregulation renders activated T cells more susceptible to cGAMP toxicity than resting T cells. In addition, we discovered that the SLC7A1 binding sites for arginine and cGAMP are distinct and that cGAMP transport can be perturbed without impacting arginine transport. Together, our study demonstrates that SLC7A1 is a potential target for small molecule drug development and gene editing for sparing tumor infiltrating T cells from collateral damage of therapeutic STING agonists or ionizing radiation, which is crucial for the curative effect of these therapies.

## Results

### Human T cell killing by extracellular cGAMP is not mediated by the known transporters

Although cGAMP and its analogs kill mouse T cells, given the clinical implications, we first wanted to test the response of primary human T cells (**Figure 1A, Supplementary Figure 1A**). Indeed, cGAMP weakly killed activated primary human CD3^+^ T cells from two unique donors with an IC_50_ of ∼20-30 µM (**Figure 1B**). 2’3’-cG^S^A^S^MP and 2’3’-CDA^S^ were both more potent than the natural cGAMP second messenger, each displaying IC_50_ values of ∼2-5 µM – even though all three molecules share similar binding affinities for STING^13,14^. We also tested several bacterial cyclic dinucleotides, which have been shown to activate human STING with weaker affinity^26,27^, in cells from one of the two donors. Surprisingly, we found that 3’3’-cGAMP kills human T cells with a comparable IC_50_ value to 2’3’-cGAMP, whereas 3’3’-cyclic di-AMP (CDA) and 3’3’-cyclic di-GMP (CDG) did not lead to any appreciable death at the highest tested concentration. Taken together, we hypothesize that differential import efficiencies of each compound may help explain the incongruence between its reported affinity for STING and toxicity toward T cells^17,23^.

**Figure 1.**
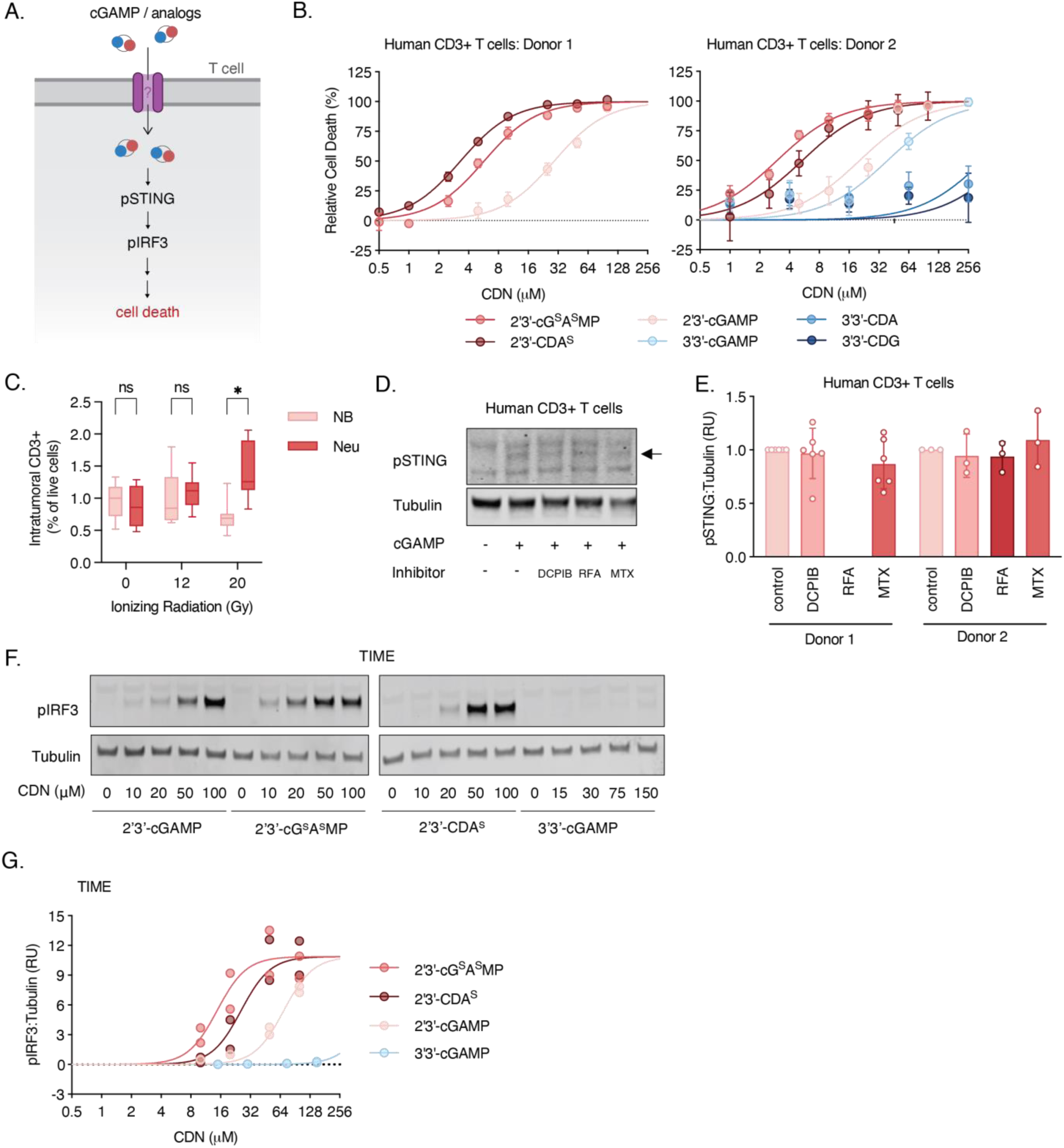
An unidentified transporter mediates human T cell killing by extracellular cGAMP. (A) Overview of cGAMP signaling in T cells. (B) Activated primary human T cells from 2 healthy donors were incubated with increasing concentrations of cyclic dinucleotides (CDNs): 2’3’-cGAMP, 2’3’-cG^S^A^S^MP, 2’3’-CDA^S^, 3’3’-cGAMP, 3’3’-CDA, and 3’3’-CDG, for 24 h in the presence of ENPP1 inhibitor, STF-1623. Relative viability was measured using Cell Titer Glo Assay (n = 3-4 biological replicates). Data are shown as mean +/- SD. (C) BALB/c mice were injected with 50,000 4T1-Luciferase cells into the mammary fat pad. Once the tumors reached 100 mm3, the tumors were irradiated with 0, 12 or 20 Gy, and injected with nonbinding (NB) or neutralizing (Neu) STING 24 h later. The mice were euthanized the following day, the tumors were extracted and CD3^+^ cells as a percentage of live cells were measured by flow cytometry. Significance are calculated by unpaired, two-tailed t-test. *p < 0.05. (D and E) Activated human primary T cells from the same 2 donors in (C) were treated with 50 µM cGAMP for 2 h, and with or without 20 µM DCPIB, 0.5 mM methotrexate (MTX), or 0.5 mM reduced folic acid (RFA), and and signaling was assessed by Western blot (n = 3-6 biological replicates). One representative Western blot from donor 2 is shown in (D) and quantification of multiple blots are shown in (E). Data are shown as the mean +/- SD. (F and G) TIME cells were treated with increasing concentration of 2’3’-cGAMP, 2’3’-cG^S^A^S^MP or 2’3’-CDA^S^ for 2 h, and signaling was assessed by Western blot (n = 2 biological replicates). One representative Western blot is shown in (F) and quantification of multiple blots is shown in (G).

We have previously shown that ionizing radiation induces extracellular cGAMP production which enters tumor infiltrating myeloid cells to activate their STING signaling^6,9^. Even though cGAMP is ∼10 times less potent at killing T cells than its clinical analogs, we next tested whether clinically relevant doses of ionizing radiation could produce enough extracellular cGAMP to kill infiltrating T cells. Indeed, we observed dose-dependent killing of tumor-infiltrating T cells when we treated mammary tumor-bearing mice with ionizing radiation at 20 Gy, but not 8 Gy (**Figure 1C)**. To specifically study the role of extracellular cGAMP in this process, we used a cell-impermeable neutralizing STING agent (Neu) to sequester extracellular cGAMP, or its non-binding mutant as a control (NB) ^6,9^. Strikingly, when we depleted extracellular cGAMP in irradiated tumors by intratumoral injection of Neu, we observed increased tumor-infiltrating T cell numbers. This increased T cell infiltration surpassed the level we observed in non-irradiated tumors injected with NB, indicating that extracellular cGAMP generated in response to high dosage ionizing radiation attracts, but also kills, T cells. These two opposing effects of endogenous extracellular cGAMP at high concentrations mirrored previous reports with the non-hydrolyzable cGAMP analog 2’3’-CDA^S^.^17,23^

We next investigated whether primary human T cells transport cGAMP using any of the previously identified transporters: LRRC8A:C/E channels used by endothelial cells^8^, fibroblasts^28^, and mouse T cells^20^, or the myeloid cell transporters SLC19A1 and SLC46A1-3^9,29^. According to published human mRNA expression data^30^, activated primary human T cells express high levels of the LRRC8A:C channel, followed by low to moderate levels of SLC19A1, SLC46A3 and SLC46A1 (**Supplementary Figure 1B**). We treated primary human T cells with inhibitors of these known transporters: methotrexate (MTX) and reduced folic acid (RFA) for SLC19A1, Sulfasalazine (SSZ) as a non-specific inhibitor for SLC19A1 and SLC46A1-3, and DCPIB as an inhibitor for LRRC8A:C/E channels. While SSZ exhibited high toxicity that precluded further analysis, none of the remaining inhibitors reduced phosphorylation of STING in primary human (**Figure 1D-E, Supplementary Figure 1C**) or mouse (**Supplementary Figure 1D-G**) T cells. DCPIB is known to be less effective in the presence of serum, so we also tested it in serum free media and observed no inhibitory effect (**Supplementary Figure 1H-I)**, suggesting that both human and mouse primary T cells use a previously unidentified transporter to import cGAMP.

To further test this hypothesis, we performed a structure-activity relationship (SAR) analysis of cGAMP and its analogs in a different cell type with known cGAMP transporters. We previously reported that TIME cells, a telomerase-immortalized human microvascular epithelial cell (HMVEC) cell line, use LRRC8A:C as its dominant transporter^8^. SAR of four cGAMP analogs and their cellular toxicity across TIME cells, and primary human T cells showed that the LRRC8A:C-dependent cell line exhibited very similar SAR patterns (**Figure 1F-G)**, while primary human T cells showed a distinct signature (**Figure 1B**). In TIME cells, 2’3’-cG^S^A^S^MP is >2-fold more potent than 2’3’-CDA^S^. However, in primary human T cells, 2’3’-CDA^S^ and 2’3’-cG^S^A^S^MP exhibited similar levels of toxicity. Consistent with our findings from the inhibitor screen, this result suggests that LRRC8A:C is not the main transporter in primary CD3^+^ T cells, and that the cGAMP transporter in human T cells is still at large.

### CRISPR screen identifies SLC7A1 as a positive regulator of extracellular cGAMP signaling in T cells

To identify the unknown T cell transporter, we aimed to conduct a genome-wide CRISPR knockout screen. First, to verify a screen-compatible model system, we tested whether the immortal human Jurkat CD4^+^ T cell line is also susceptible to cGAMP toxicity and uses the same transporter(s) as primary human T cells. Mirroring our results in primary human CD4^+^ T cells, none of the transporter inhibitors had significant effects on STING signaling in either resting Jurkat (**Supplementary Figure 2A-B**) or Jurkat activated with PMA/ionomycin (**Figure 2A-B**). Furthermore, SAR of cGAMP analogs in both resting and activated Jurkat T cells are similar to that observed with primary human T cells (**Figure 2C-D, Supplementary Figure 2C-D**). Therefore, we proceeded to use Jurkat as a model cell line for a genome-wide survival screen (**Figure 2E**), reasoning that knockout of factors involved in cGAMP-mediated cell death – including the cGAMP transporter – would allow a cell to survive treatment with a lethal dose of cGAMP. We treated a pool of 200 million cells carrying the 200,000 sgRNA library (10 guides per gene) with a LD_30_ dose of cGAMP, a concentration that kills 30% of the cells every day. Following 10 days of treatment, we measured sgRNA enrichment among the surviving population to identify genes contributing to cGAMP-mediated toxicity.

**Figure 2.**
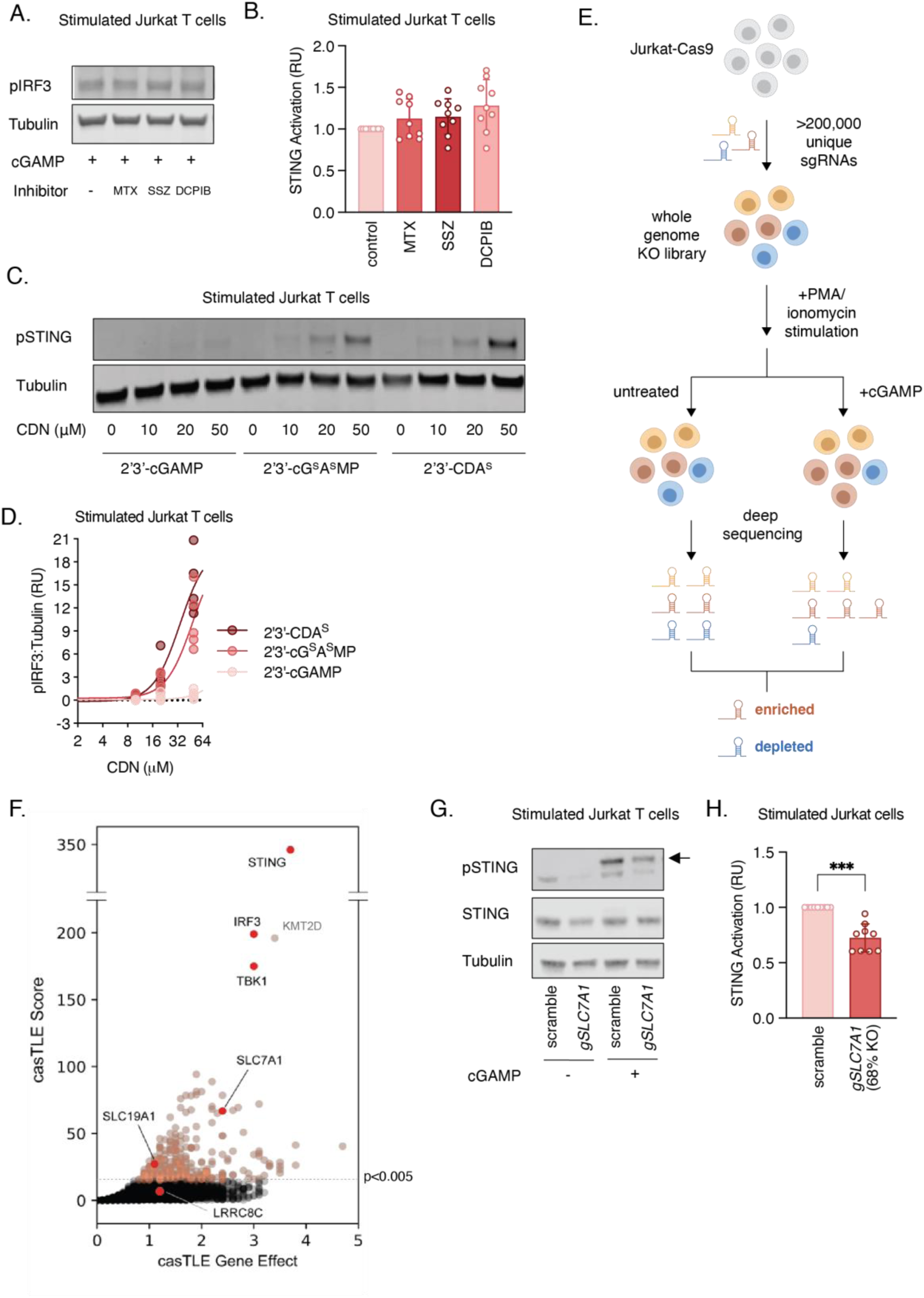
CRISPR screen identifies SLC7A1 as a positive regulator of extracellular cGAMP signaling in T cells. (A and B) Jurkat cells were treated with 5 ng/mL PMA and 500 ng/mL ionomycin for 6 hrs. 72 hours later, stimulated cells were treated with 50 µM cGAMP for 2 h, and with or without 0.5 mM methotrexate (MTX), 1 mM sulfasalazine (SSZ) or 20 µM DCPIB, and STING activation (pIRF3:Tubulin or pSTING:Tubulin) was assessed by Western blot (n = 9 biological replicates). One representative Western blot is shown in (A) and quantification of multiple blots is shown in (B). Data are shown as the mean +/- SD. (C and D) Jurkat cells were treated with 5 ng/mL PMA and 500 ng/mL ionomycin for 6 hrs. 72 hours later, stimulated cells were treated with increasing concentration of 2’3’-cGAMP, 2’3’- cG^S^A^S^MP or 2’3’-CDA^S^ for 2 h, and signaling was assessed by Western blot (n = 5 biological replicates). One representative Western blot is shown in (C) and quantification of multiple blots is shown in (D). Data are shown as the mean +/- SD. (E) CRISPR screen overview. Whole genome-targeting sgRNAs were introduced lentivirally into a Jurkat-Cas9 cell line, which was then treated with PMA and ionomycin. 24 hrs later, stimulated library cells were treated daily with LD_30_ cGAMP or untreated for 10 days. Genomic DNA was harvested, deep sequenced, and analyzed for sgRNA enrichment or depletion. (F) Plot of casTLE score with positive effect size. Known regulators of STING pathway and SLC7A1 are marked in red. A p<0.005 significance threshold is indicated as a dotted line, points exceeding this threshold are marked in orange. (G and H) Jurkat cells were transduced with Cas9 along with non-targeting (scramble) or SLC7A1-targeting sgRNA, were treated with 75 uM cGAMP for 2h, and STING activation (pIRF3:Tubulin or pSTING:Tubulin) was assessed by Western blot (n = 9 biological replicates). Average knock-out score across independent experiments was determined by ICE (Synthego) to be 68%. One representative Western blot is shown in (G) and quantification of multiple blots is shown in (H). Significance calculated by unpaired, two-tailed t-test. Data are shown as mean +/- SD. ***p < 0.001

As expected, genes encoding STING, TBK1, and IRF3 arose to have the highest statistical significance and fold enrichment (**Figure 2F**). The gene encoding solute carrier protein SLC7A1 emerged as a top candidate transporter. It has been previously characterized as a cationic amino acid transporter that transports substrates such as L-arginine^31^. Activated T cells rapidly expand and increase their metabolic consumption by upregulating SLC7A1 mRNA levels by ∼30 fold (**Supplementary Figure 2E)** and protein levels by ∼10 fold (**Supplementary Figure 2F**) to take in more arginine. SLC7A1 knockout in T cells has previously been shown to arrest cell cycle progression and diminish cytokine release^32^, and indeed we observed that Jurkat SLC7A1^-/-^ cell viability decreased after multiple passages. To circumvent this viability limitation, we measured extracellular cGAMP-mediated IRF3 phosphorylation in a heterogeneous pool of Jurkat cells immediately following SLC7A1 knockout. We achieved SLC7A1 knockout in ∼60-70% of cells, which led to a 40% reduction in pIRF3 in the resting cellular pool upon cGAMP treatment (**Supplementary Figure 2G-H**). In the same cellular pool, we observed a 30% reduction of pIRF3 in stimulated cells (**Figure 2G-H**). These results suggest that SLC7A1 plays a role in extracellular cGAMP signaling in Jurkat T cells, nominating it as a possible cGAMP transporter.

### SLC7A1 is important for cGAMP transport and STING signaling in primary human and mouse T cells

Next, we investigated whether SLC7A1 is also involved in extracellular cGAMP signaling in primary T cells. We followed the same strategy to create heterogeneous SLC7A1 knockout pools of primary human T cells from two donors, and treated the cells with cGAMP during the brief 72 hours window before significant cell death was observed from arginine starvation. Partial knockout of SLC7A1 in donor 1’s primary human CD3^+^ by 48% reduced pSTING levels by ∼25% compared to the non-targeting control guide, whereas 73% knockout of SLC7A1 in cells from donor 2 led to a ∼40% reduction of pSTING levels (**Figure 3A-B)**. Likewise, the heterogenous pool of SLC7A1 knockout mouse primary CD4^+^ cells (∼30% indel score) responded 25% less and CD8^+^ cells (∼60% indel score) responded 55% less to extracellular cGAMP-induced p-IRF3 than their WT counterparts (**Figure 3C-D**). Importantly, these differences were abolished when cGAMP was electroporated to bypass the need for transport (**Figure 3E-F**). We then performed a direct binding assay with purified flag-tagged full-length SLC7A1 and 3’3’-cGAMP photoaffinity probe (**Supplementary Figure 3A**). If SLC7A1 binds to the probes, it can be covalently crosslinked with the probes via diazirine upon UV irradiation at 365 nm and visualized in gel by conjugating the alkyne handle of the probe to fluorescence tag (R110-N_3_) via click chemistry (**Figure 3G**). Indeed, the probe successfully crosslinked with SLC7A1 in a UV crosslinking dependent manner, as evidenced by the fluorescent signal as distinct bands corresponding to the FLAG bands (**Supplementary Figure 3B**). Utilizing this probe, we performed competition experiment and observed that both 2’3’-cGAMP and 3’3’-cGAMP inhibited the probe from labeling SLC7A1 with 3’3’-cGAMP expectedly being a more potent competitor (**Figure 3H**). Together, our data demonstrates that SLC7A1 mediates STING signaling induced by extracellular cGAMP in primary T cells by facilitating cGAMP import.

**Figure 3.**
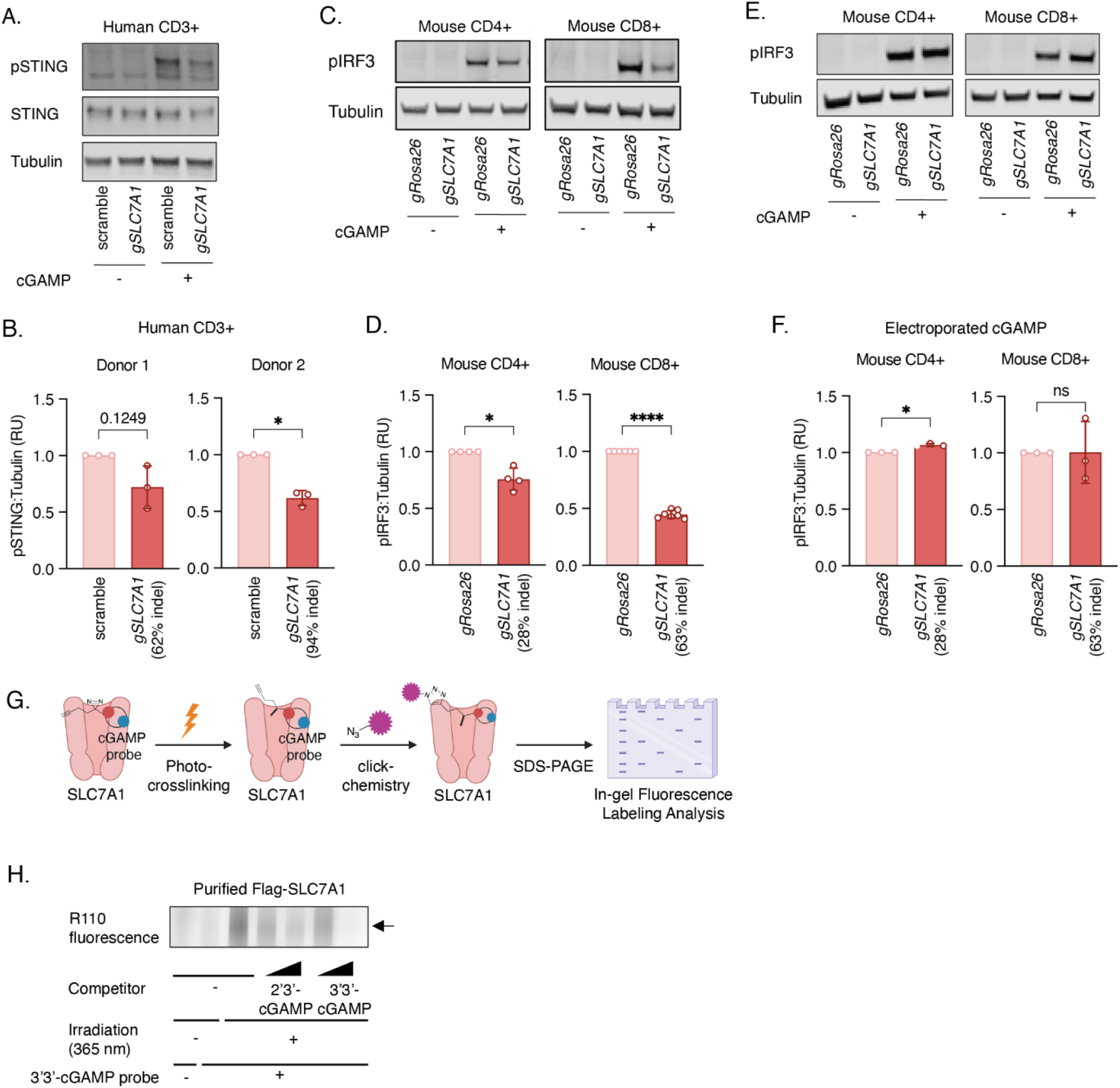
SLC7A1 is important for cGAMP transport and STING signaling in primary human and mouse T cells. (A and B) Activated human primary CD3^+^ T cells from 2 healthy donors were electroporated with Cas9 protein and non-targeting (scramble) or SLC7A1-targeting sgRNA. Cells were treated with 50 µM cGAMP for 2 h three days later, and signaling was assessed by Western blot (n = 3 biological replicates). Indel score was determined by ICE (Synthego). One representative Western blot from Donor 2 is shown in (A) and quantification of multiple blots from 2 donors are shown in (B). Significance calculated by unpaired, two-tailed t-test. Data are shown as the mean +/- SD. *p < 0.05 (C and D) Activated mouse CD4+ and CD8+ T cells were electroporated with Cas9 protein and Rosa26-targeting or SLC7A1-targeting sgRNA. Cells were treated with 50 µM cGAMP for 3 h three days later, and signaling was assessed by Western blot (n = 4 - 6 biological replicates). One representative Western blot from each group of CD4+ and CD8+ T cells are shown in (C) and quantification of multiple blots from the two groups are shown in (D). Significance calculated by unpaired, two-tailed t-test. Data are shown as the mean +/- SD. *p < 0.05, ****p<0.0001 (E and F) Rosa26- and SLC7A1-knocked out CD4+ and CD8+ mouse T cells from (C and D) were electroporated with 100 nM of cGAMP for 2 h, and signaling was assessed by Western blot (n = 3 biological replicates). One representative Western blot from each group of CD4+ and CD8+ T cells are shown in (E) and quantification of multiple blots from the two groups are shown in (F). Significance calculated by unpaired, two-tailed t-test. Data are shown as the mean +/- SD. *p < 0.05. (G) Schematic of diazirine probe crosslinking assay. Purified FLAG-SLC7A1 proteins were incubated with a 3’3’-cGAMP probe containing a diazirine motif, and then UV irradiated at 365 nm. Crosslinked proteins were then conjugated with a R110 fluorescence tag containing an alkyne handle (R110-N_3_) via click chemistry. Proteins were visualized were visualized on SDS-PAGE gel and R110 fluorescence and Flag signal was assessed by Western Blot. (H) Purified FLAG-SLC7A1 protein was incubated with 500 µM of 3’3’-cGAMP diazirine probe, and with or without 2’3’- or 3’3’-cGAMP (lower concentration: 500 µM, higher concentration: 2mM), and irradiated at 365 nm for 20 min. Crosslinked protein signals were assayed by Western Blot. Flag blot is shown in (Supplementary Figure 3B) and R110 fluorescence blot is shown in (H).

### cGAMP and arginine bind distinct sites on SLC7A1

Puzzled by the fact that SLC7A1 can transport substrates that are not only structurally distinct, but also different in their charges, we hypothesized that it might bind different substrates at different sites. Molecular modeling predicted a cGAMP binding site on the extracellular surface of SLC7A1, >16 Å away from the arginine binding site modeled in the interior^33^ (**Figure 4A**). To experimentally test the molecular model, we performed mutagenesis studies in a doxycycline-inducible heterologous SLC7A1 expression system. SLC7A1 can only be lowly expressed when introduced exogenously into U937 cells, which yielded a small but significant decrease in STING signaling and cell viability 24h after induction (**Figure 4B, Supplementary Figure 4A, B**). These small increases are not due to expression changes in the STING pathway components or other cGAMP transporters as expressions of these genes are unaltered upon doxycycline-induction (**Supplementary Figure 4C**). As expected, SLC7A1 expression also increased [^3^H] L-arginine intake in these cells (**Supplementary Figure 4D**) which also basally uptake arginine (**Supplementary Figure 4E**). By introducing point mutations in this exogenous SLC7A1 expression system, we observed that the predicted arginine binding site mutation (S354M) completely abolished arginine transport without affecting cGAMP-mediated cell killing (**Figure 4B-C**). We then performed single and double mutations around the predicted cGAMP binding site and observed that R59A/D404A double mutant and R130A single mutant all decreased cGAMP-mediated cell killing without affecting arginine transport (Supplementary Figure 4F-G). The R59A/R130A double mutant and D404A/R130A double mutant also decreased cGAMP-mediated cell killing without affecting arginine transport. Remarkably, the R59A/D404A/R130A triple mutant completely abolished the effect of 20 μM extracellular cGAMP without affecting arginine transport **(Figure 4B-C)**. The identification of the triple mutant triangulated the cGAMP binding pocket on SLC7A1 and validated our molecular modeling results (**Figure 4D**). Interestingly, while the triple mutant also abolished 2’3’-CDA^s^ and 2’3’-cG^s^A^s^MP induced cell death via SLC7A1, it did not completely abolish that of ‘3’3-cGAMP, suggesting that 3’3’-cGAMP engages with additional residues explaining why it is a better substrate (**Figure 4E**). Finally, cGAMP does not compete with direct arginine uptake by SLC7A1, while lysine does (**Figure 4F**). Together, these biochemical assays definitively demonstrated that SLC7A1 is a cGAMP transporter and uses separate binding sites for different substrates.

**Figure 4.**
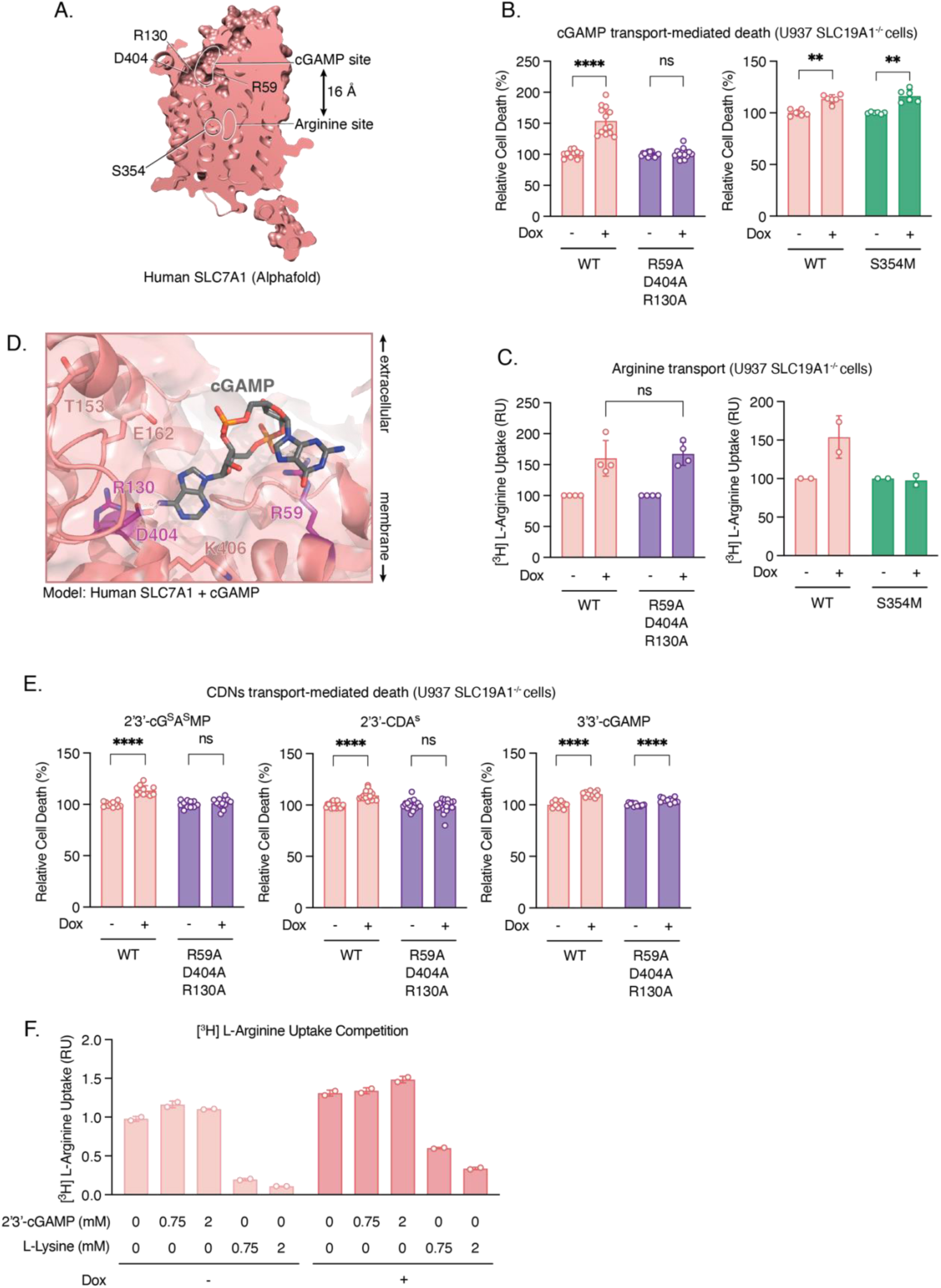
cGAMP and arginine bind distinct sites on SLC7A1. (A) AlphaFold-predicted structure of human SLC7A1. The modeled cGAMP and Arginine binding sites are outlined in white. (B) U937 SLC19A1^-/-^ -tet-SLC7A1 (WT, R59A/D404A/R130A or S354M)-Flag cells were incubated with or without 1 µg/mL doxycycline (dox) for 24 h. Cells were then treated with 20 µM 2’3’-cGAMP for 24 h in the presence of ENPP1 inhibitor, STF-1623. Relative viability was measured using Cell Titer Glo Assay (n = 6-14 biological replicates, as indicated). Significance calculated by unpaired, two-tailed t-test. Data are shown as the mean +/- SD. **p<0.01, ****p<0.0001 (C) U937 SLC19A1^-/-^ -tet-SLC7A1 (WT, R59A/D404A/R130A or S354M)-Flag cells were incubated with or without 1 µg/mL doxycycline (dox) for 24 h. Cells were then treated with 21 nM ^3^H-Arginine for 1 min (n = 2-4 biological replicates, as indicated), and radioactivity was measured using a scintillation counter and normalized to protein amount. (D) AlphaFold-predicted structure of human SLC7A1 with cGAMP (gray sticks) docked into a pocket on the extracellular-facing surface. Residues mutated are shown as sticks; residues validated to have an effect on cGAMP transport shown in purple. Residues 229-251 (a flexible loop with low AlphaFold prediction score) are omitted for ease of visualization. (E) U937 SLC19A1^-/-^ -tet-SLC7A1 (WT or R59A/D404A/R130A)-Flag cells were incubated with or without 1 µg/mL doxycycline (dox) for 24 h. Cells were then treated with 2 µM 2’3’-cG^s^A^s^MP, 4 µM 2’3’-CDA^s^ or 40 µM 3’3’-cGAMP for 24 h in the presence of ENPP1 inhibitor, STF-1623. Relative viability was measured using Cell Titer Glo Assay (n = 11-19 biological replicates, as indicated). Significance calculated by unpaired, two-tailed t-test. Data are shown as the mean +/- SD. ****p<0.0001 (F) U937 SLC19A1^-/-^ -tet-SLC7A1-Flag cells were incubated with or without 1 µg/mL doxycycline (dox) for 24 h. Cells were then treated with 21 nM ^3^H-Arginine and different concentrations of 2’3’-cGAMP or L-Lysine for 1 min (n = 2 biological replicate), and radioactivity was measured using a scintillation counter and normalized to protein amount.

### Homolog SLC7A2 also transports cGAMP

SLC7A1 is a member of the larger SLC7 family, of which SLC7A2 is the most closely related: sharing 58% sequence identity and 71% sequence similarity with SLC7A1. SLC7A2 is also a cationic amino acid transporter, but has a different cell type expression profile and is notably absent from immune cells (**Supplementary Figure 5A**). Although SLC7A2 is not expressed by T cells, we reasoned that it might also transport cGAMP given its close homology to SLC7A1. To test this hypothesis, we set up an analogous dox-inducible, FLAG-tagged system for heterologous SLC7A2 expression in *SLC19A1*^-/-^ U937 cells. Dox induction led to robust protein expression – significantly higher than the expression we achieved with SLC7A1 (**Supplementary Figure 4A**) – with smearing consistent with its predicted glycosylation (**Figure 5A**). A large increase in ^3^H-arginine uptake after SLC7A2 overexpression confirmed its functionality in this heterologous system (**Supplementary Figure 5B**). SLC7A2 overexpression greatly increased extracellular cGAMP-mediated pIRF3 signaling (**Figure 5A-B**) but had no effect when the need for cGAMP transport was bypassed by electroporating cGAMP directly into cells (**Figure 5C-D**). This increase in pIRF3 was not due to any effect of altered amino acid uptake, as we saw a similar increase in pIRF3 when we cultured cells in media lacking the SLC7A2 amino acid substrates arginine, lysine, histidine, and ornithine (**Supplementary Figure 5C-D**). This increase in pIRF3 was also not due to expression changes in the STING pathway components or other cGAMP transporters as expressions of these genes are unaltered upon doxycycline-induction (**Supplementary Figure 5E**). Similar to SLC7A1, overexpression of SLC7A2 drastically shifted the dose response curve of cGAMP-mediated U937 cell killing (**Figure 5E**). Most importantly, overexpression of SLC7A2 directly increased cGAMP transport into the cells (**Figure 5F**). Molecular modeling predicted a cGAMP binding site in SLC7A2 similar to that in SLC7A1, far away from the Arginine binding site (**Figure 5G**). Our mutagenesis studies showed that K61A and D406A mutant each individually abolished cGAMP transport activity without abolishing arginine transport activity (**Figure 5H-J)**. Interestingly, the E234A mutant increased cGAMP transport without affecting arginine transport, likely because the glutamate at 234 position is repulsing the electrons in the guanine ring. Finally, the S356M mutant abolished arginine transport activity without affecting cGAMP transport demonstrating that SLC7A2 also has a separate binding pocket for cGAMP than for arginine. Together, our data suggest that SLC7A2, like SLC7A1, is a cGAMP transporter. But unlike SLC7A1 which is highly expressed on activated T cells, SLC7A2 has a broad expression profile across different tissues and cancer types. Future studies are warranted to explore the physiological relevance of SLC7A2’s cGAMP transport activity.

**Figure 5.**
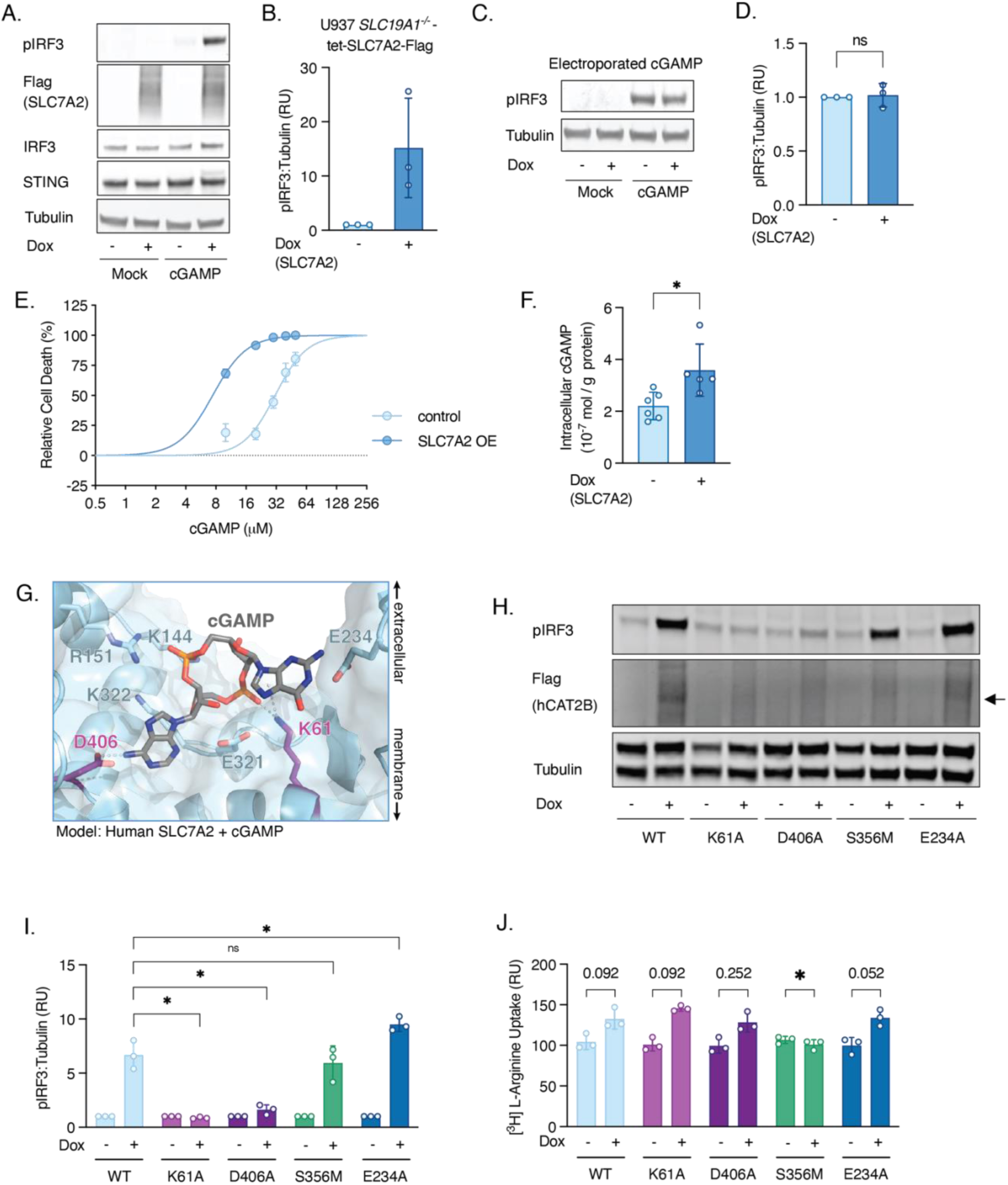
SLC7A2, a close homolog of SLC7A1, also imports cGAMP. (A and B) U937 SLC19A1^-/-^ -tet-SLC7A2-Flag cells were incubated with or without 1 µg/mL doxycycline (dox) for 24 h. Cells were then treated with 50 µM cGAMP for 2 h, and signaling was assessed by Western blot (n = 3 biological replicates). One representative Western blot is shown in (A) and quantification of multiple blots is shown in (B). Data are shown as mean +/- SD. (C and D) U937 SLC19A1^-/-^ -tet-SLC7A2-Flag cells were incubated with or without 1 µg/mL doxycycline (dox) for 24 h. Cells were then electroporated with 100 nM cGAMP for 2 h, and signaling was assessed by Western blot (n = 3 biological replicates). One representative Western blot is shown in (C) and quantification of multiple blots is shown in (D). Significance calculated by unpaired, two-tailed t-test. Data are shown as mean +/- SD. (E) U937 SLC19A1^-/-^ -tet-SLC7A2-Flag cells were incubated with or without 1 µg/mL doxycycline (dox) for 24 h. Cells were then incubated with increasing concentrations of 2’3’-cGAMP for 24 h in the presence of ENPP1 inhibitor, STF-1623. Relative cell count was measured using Attune NxT Flow Cytometer (n = 4 biological replicates). Data are shown as the mean +/- SD. (F) U937 SLC19A1^-/-^ -tet-SLC7A2-Flag cells were incubated with or without 1 µg/mL doxycycline (dox) for 24 h. Cells were then incubated with 10 µM cGAMP at 37°C for 15 min in the presence of ENPP1 inhibitor STF-1623 (n = 5-6 biological duplicates, as indicated. One extreme outlier with value 29.8 10^-7^ mol/g protein, from the dox-treated group was omitted after performing Grubb’s test). Cells were then washed in cold PBS, lysed in T-PER lysis buffer, and intracellular cGAMP level was determined using cGAMP ELISA kit and normalized to protein amount. Significance calculated by unpaired, two-tailed t-test. Data are shown as the mean +/- SD. *p<0.05. (G) AlphaFold-predicted structure of human SLC7A2 with cGAMP docked into a pocket on the extracellular-facing surface. Residues mutated are shown as sticks; residues validated to have an effect on cGAMP transport shown in purple. (H and I) U937 SLC19A1^-/-^ -tet-SLC7A2 (WT, E234A, S356M, K61A or D406A)- Flag cells were incubated with or without doxycycline (dox) for 24 h. Cells were then treated with 50 µM cGAMP, and signaling was assessed by Western blot (n = 3 biological replicates). One representative Western Blot is shown in (H), and quantification of multiple blots is shown in (I). Significance calculated by unpaired, two-tailed t-test. Data are shown as mean +/- SD. *p<0.05 (J) U937 SLC19A1^-/-^ -tet-SLC7A2 (WT, E234A, S356M, K61A or D406A)- Flag cells were incubated with or without doxycycline (dox) for 24 h. Cells were then treated with 21 nM ^3^H-Arginine for 1 min (n = 3 biological replicates), and radioactivity was measured using a scintillation counter and normalized to protein amount. Significance calculated by unpaired, two-tailed t-test. Data are shown as mean +/- SD.

## Discussion

In this study, we identified SLC7A1 as the highly sought after transporter responsible for STING agonist-induced T cell toxicity. Our screening approach, modified from our previous work^8,29^, succeeded in identifying the T cell transporter because we first identified a model cell line, Jurkat T cells, which share the same cyclic dinucleotide substrate selectivity and import inhibitor profiles as primary human T cells. In addition, Jurkat T cells can be activated like primary T cells and, when activated, are also susceptible to toxicity caused by cGAMP and its analogs. Our previous whole genome CRISPR screens in U937 cells identified SLC19A1 as the first cGAMP transporter^29^ and LRRC8A:C and LRRC8A:E channels as the cGAMP transporters in endothelial cells and fibroblasts respectively^8^, but did not identify SLC7A1 due to its lower expression level in U937 cells.

Previous work reported that LRRC8C is a cGAMP transporter in mouse T cells and mediates mouse T cell death by cGAMP-STING-induced p53 activation^20^. Since we previously observed differences between mouse and human transporters in their ability to import cGAMP^9,29^, we prioritized testing the relative importance of SLC7A1 and LRRC8A:C channels in activated primary T cells from both species. We conclude that SLC7A1 is the dominant T cell transporter in both mouse and human T cells, given that knockout of SLC7A1 impaired response to imported cGAMP similarly in both mouse and human T cells (Figure 3). Two factors likely contribute to SLC7A1’s functional dominance compared to the LRRC8A:C channel in activated primary T cells. First, SLC7A1 is upregulated to a level much higher than that of the LRRC8A:C channel when T cells are activated and start proliferating due to their increased demand for cationic amino acids. Second, SLC7A1 is a solute carrier, and it transports cGAMP down its electrochemical gradient. When cGAMP analogs are used therapeutically, the extracellular concentrations are much higher than intracellular concentrations. The LRRC8A:C channel, however, is only somewhat active at isotonic conditions. The pore is opened by hypotonicity, which occurs less frequently in T cells than in endothelial cells where it is the dominant cGAMP transporter^8^.

With the addition of SLC7A1, the complexity and diversity of cGAMP transporters continues to grow and suggests a finely tuned response mechanism to cGAMP when it is secreted as an immunotransmitter. The biological importance of each cGAMP transporter depends on its relative expression levels on the cell type of interest, its cGAMP transport efficiency, and its function transporting other substrates. SLC7A1’s cGAMP transport activity is weak compared to its arginine transport activity, so its upregulation in primary T cells has a concentration window where it promotes T cell proliferation without cGAMP toxicity. A microenvironment is a mixture of stromal cells, myeloid cells, and T cells. We hypothesize that the strong cGAMP transporters in stromal cells and myeloid cells ensure that virally induced or cancer secreted cGAMP activates STING in the stroma and myeloid cells to produce type-I interferon and mount potent anti-viral and anti-cancer immunity. This is because compared to SLC7A1 which is lowly expressed in naive T cells, SLC46A2 is highly expressed by primary human monocytes and transports cGAMP more efficiently in these cells; the LRRC8A:C complex is highly expressed by endothelial cells and LRRC8A:E complex is highly expressed by fibroblasts and both transport cGAMP efficiently basally but much more so with cell swelling^8,10,34,35^. However, we hypothesize that the increase of SLC7A1 expression in T cells as they become activated with rapid proliferation allows cGAMP-mediated T cell killing once they have performed their effector functions to avoid autoimmunity. In a way, SLC7A1’s cGAMP transport activity in T cells could function as an adaptive immune checkpoint. Future studies are warranted to test these hypotheses.

In the case of cancer immunotherapy, blocking this adaptive immune checkpoint could potentially prolong the lifespan of activated T cells. This could be used during treatments with exogenous cGAMP or cGAMP analogs, or when high concentrations of endogenous extracellular cGAMP are generated after high dose ionizing radiation. Since arginine and cGAMP transport by SLC7A1 are functionally separable, specifically inhibiting cGAMP but not arginine transport would be a viable strategy to block only cGAMP analog-mediated T cell toxicity without affecting T cell metabolism. This would also permit STING activation in other cell types essential for antiviral and anti-cancer immunity. SLC7A1 is the first cGAMP transporter that is an actionable drug target for boosting the efficacy of cancer immunotherapy and other treatment mechanisms that rely on T cell functions.

In addition, we show that SLC7A1’s homolog SLC7A2, which is not expressed in T cells, also imports cGAMP in a manner that is separable from arginine import. SLC7A2 is highly expressed in cancer cells and its physiological role there needs to be explored. Since these transporters are conserved in sequence and molecular function, it would be important to consider the impact of any potential inhibitors on both transporters, including other therapeutic contexts for targeting SLC7A2 or undesired consequences of inhibiting both transporters.

## Acknowledgments

We thank A. Pawluk, B. Plosky, C. Ricci-Tam, and the Arc Institute Scientific Publications Team for constructive feedback on the manuscript. We thank C. Ritchie for generating U937 SLC19A1^-/-^ cell line. We thank Gaelen Hess and the Bassik Lab (Stanford Genetics) for providing the whole genome CRISPR library plasmid and use of NextSeq 500 sequencer. We thank all Li Lab members for their constructive comments and discussion through the course of this study. V.S. was supported by Stanford Graduate Fellowship. D.R.C. was supported by Stanford Graduate Fellowship and Sarafan ChEM-H Chemistry/Biology Interface Training Program. J.M.L. was supported by Stanford Knight-Hennessy Scholarship and Sarafan ChEM-H Chemistry/Biology Interface Training Program. A.C. was supported by NIH 5T32GM736544 and 1F30CA250145. This work was supported by NIH DP2CA228044 (L.L)., NIH 5R01CA258427-02 (L.L.)., and Arc Institute.

## Author contributions

V.S., D.R.C, J.M.L., and L.L. designed research; V.S., D.R.C., J.M.L. J.A.C., X.C., and A.F.C. performed research; V.S., D.R.C., and J.M.L. contributed new reagents/analytic tools; V.S., D.R.C., J.M.L., and A.F.C. analyzed data; and V.S., J.A.C., and L.L. wrote the paper.

## Materials and Methods

### Methods

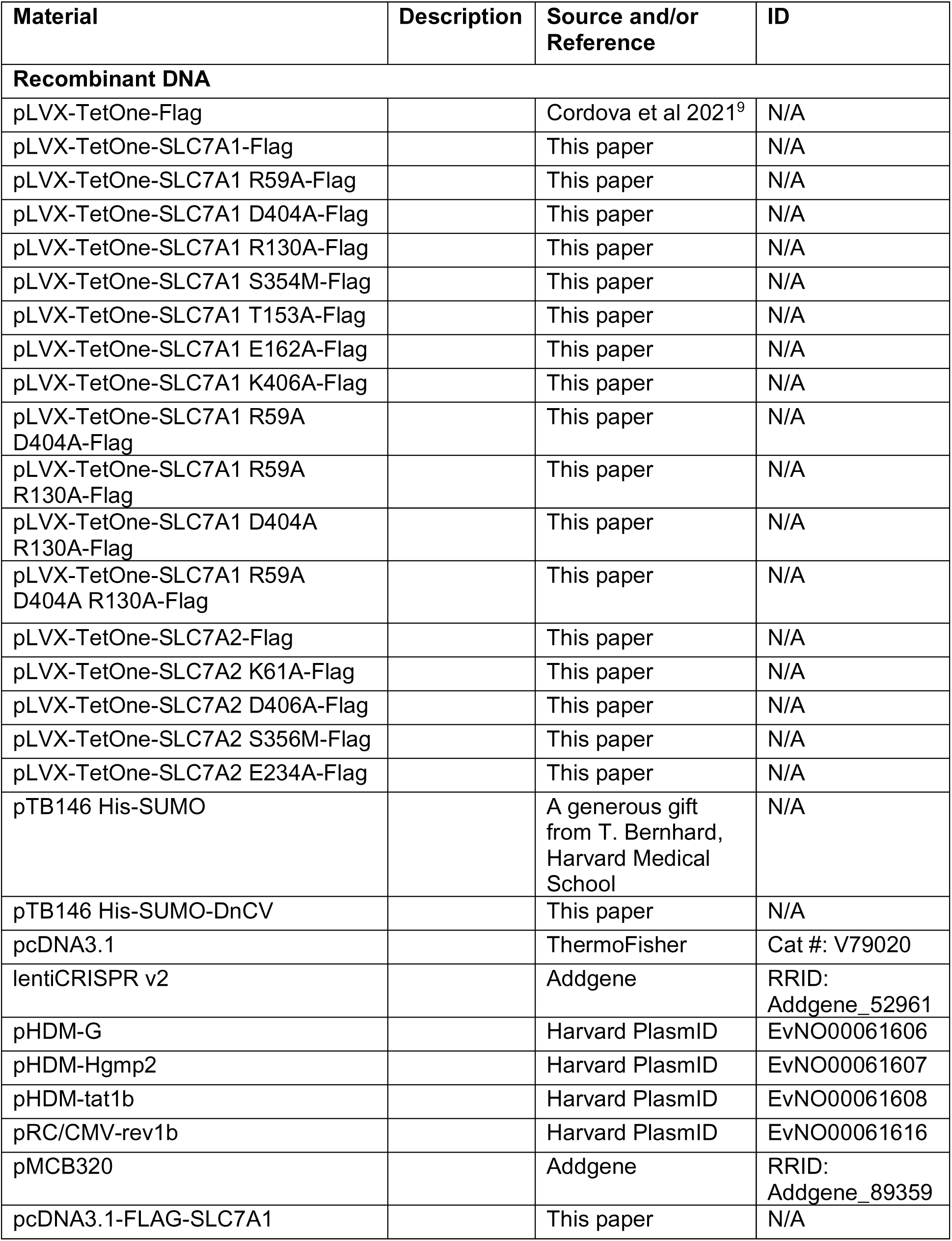
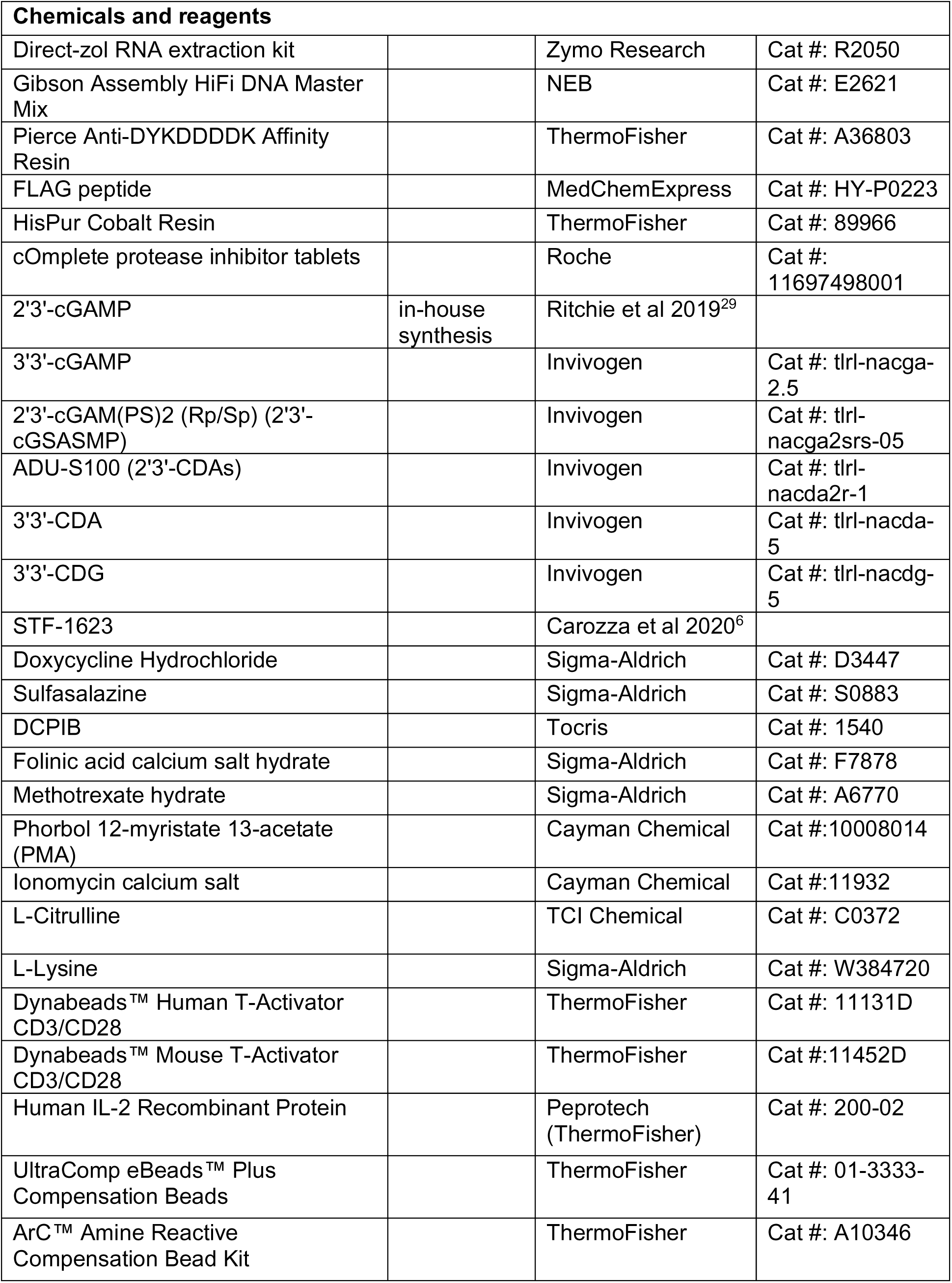
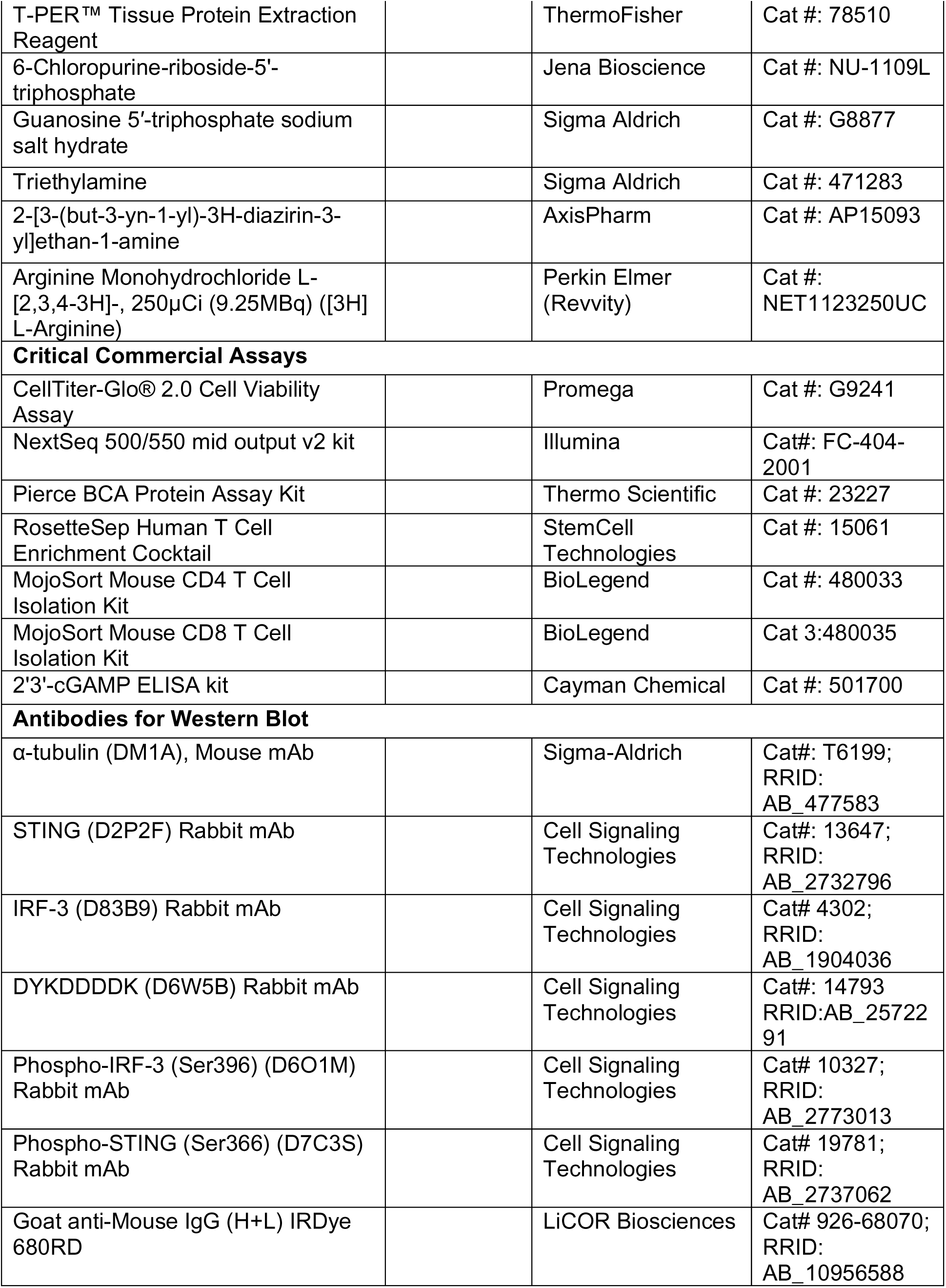
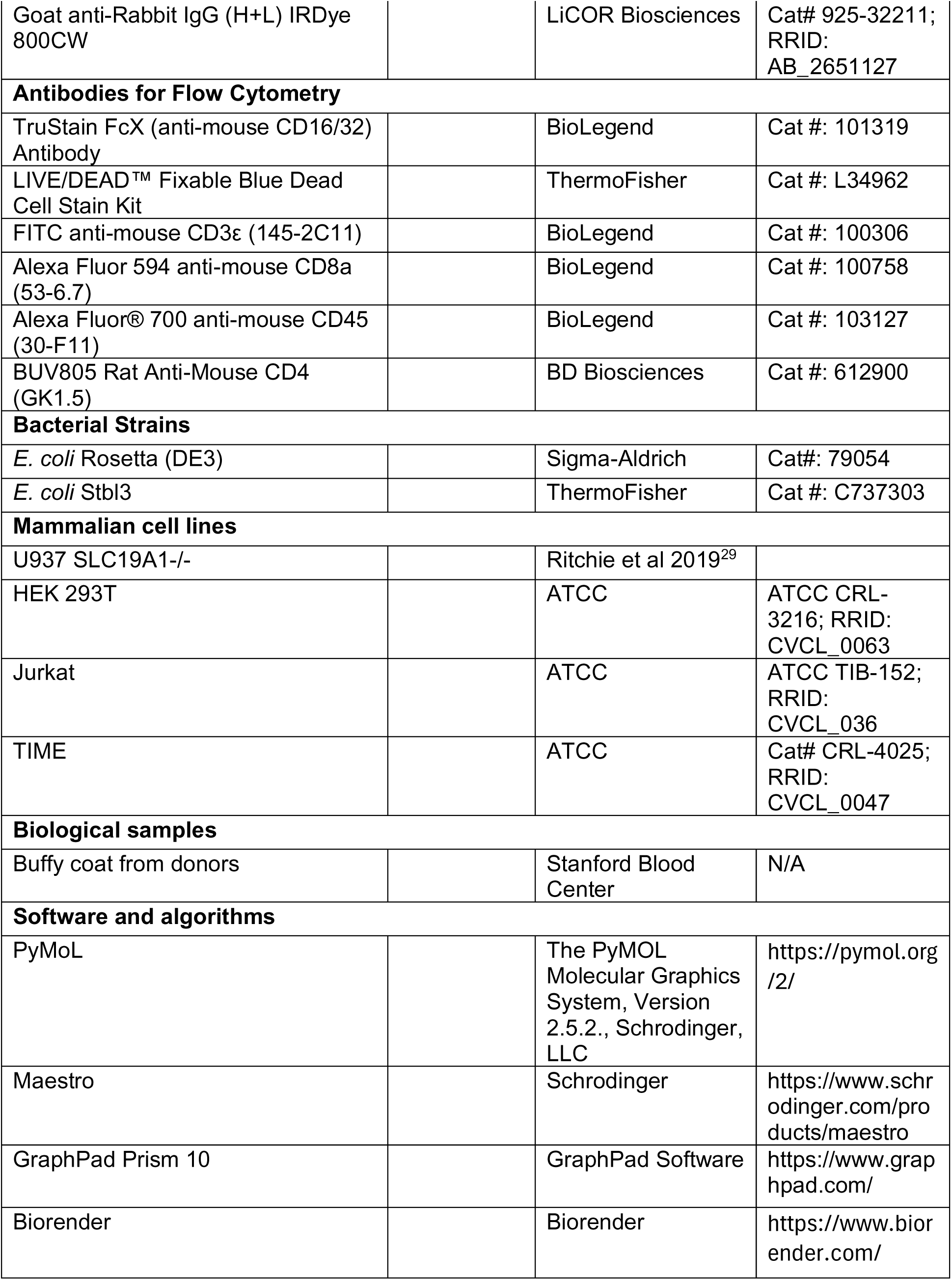
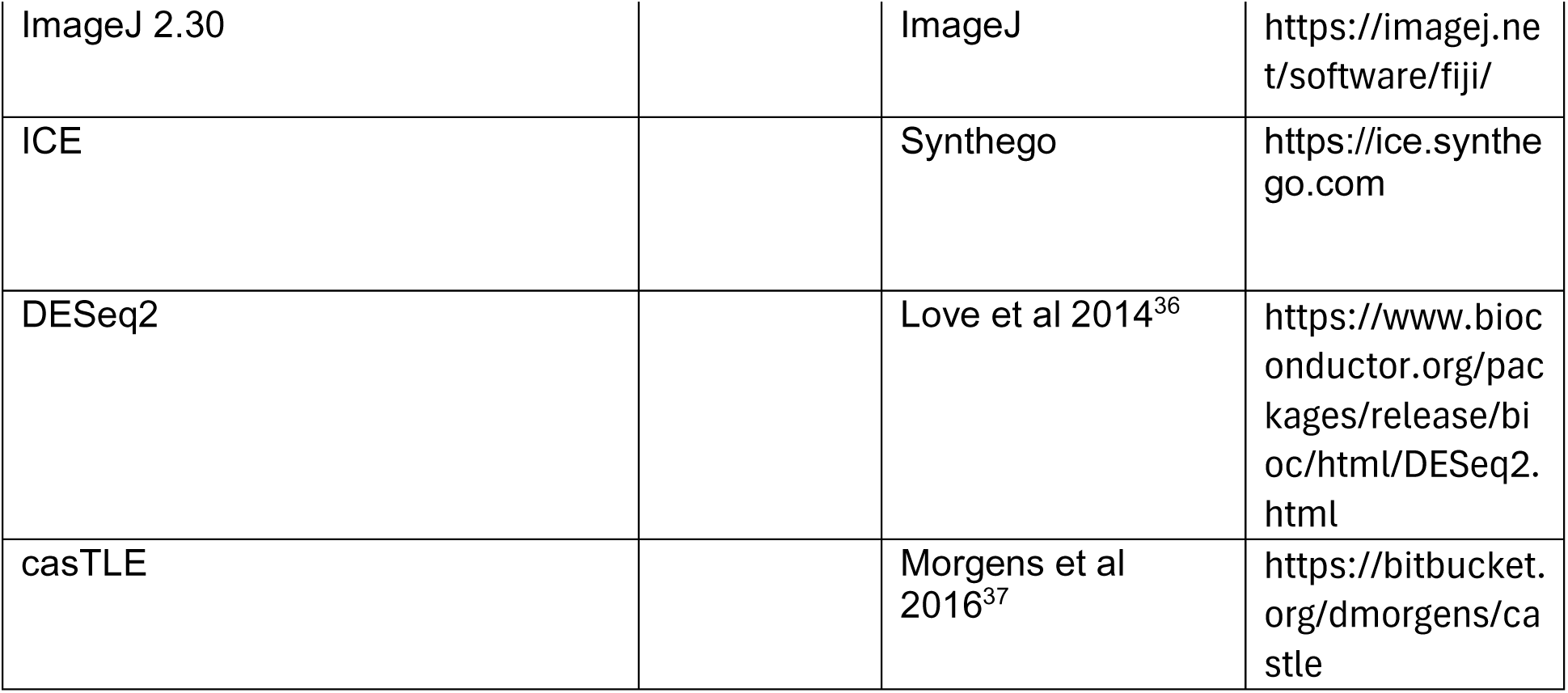

#### Cell Culture

HEK 293T cells used for lentivirus generation were maintained in DMEM with L-glutamine, 4.5 g/L glucose, and sodium pyruvate (Corning) supplemented with 10% FBS (Atlanta Biologicals) and 1% penicillin-streptomycin (GIBCO). Telomerase-immortalized human microvascular endothelial (TIME) cells were maintained in vascular cell basal media supplemented with microvasculature endothelial cell growth kit-VEGF (ATCC) and 0.1% penicillin-streptomycin (GIBCO). U937 cells, Jurkat cells and Human CD3^+^ primary T cells were maintained in RPMI (Corning) supplemented with 10% heat-inactivated FBS (Atlanta Biologicals) and 1% penicillin-streptomycin (GIBCO). Mouse CD4+ and CD8+ primary T cells were maintained in RPMI (Corning) supplemented with 10% heat-inactivated FBS (Atlanta Biologicals), 1% penicillin-streptomycin (GIBCO), and 50 µM 2-Mercaptoethanol (Sigma-Aldrich). Both human and mouse T cells were also supplemented with 300 U/mL rIL-2 (PeproTech). Cells were maintained in a 5% CO_2_ incubator at 37°C. Expi293 cells were maintained in FreeStyle medium (Gibco) supplemented with 33% Expi293 Expression medium (Gibco) in an 8% CO_2_ incubator at 37°C.

#### Recombinant DNA

A plasmid containing the CDS of human SLC7A2 (pDONR221_SLC7A2) was purchased from Addgene. A custom plasmid (pTwist-CMV) containing the coding sequences of human SLC7A1 was purchased from Twist Bioscience. A custom plasmid (pTB146 His-SUMO-DncV) containing full-length *Vibrio cholerae* DncV cDNA sequence^38^ was purchased from Twist Bioscience.

To generate doxycycline inducible lentiviral plasmids, the transporter CDS was amplified from the appropriate plasmid and cloned into an EcoRI/ BamHI linearized pLVX-TetOne-FLAG-Blast plasmid by isothermal Gibson assembly. To introduce point mutations in SLC7A1, the CDS was amplified with primers carrying the corresponding base substitutions and cloned into linearized pLVX-TetOne-FLAG-Blast plasmid by isothermal Gibson assembly. To generate plasmid for SLC7A1 protein expression, the SLC7A1 CDS was amplified with an N-terminal FLAG tag and a HRV-3C protease sequence linker, and cloned into pcDNA3.1 plasmid (ThermoFisher) by isothermal Gibson Assembly. Primers used for cloning can be made available upon request.

##### Reagents and antibodies

2’3’-cyclic-GMP-AMP (cGAMP) and soluble STING were synthesized and purified in-house as previously described^29^. 2’3’-bisphosphorothioate-cyclic-GMP-AMP (2’3’-cG^S^A^S^MP), 2’3’-cyclic-di-AMP (2’3’-CDA), 2’3’-bisphosphorothioate-cyclic-diAMP (2’3’-CDA^S^), 3’3’-cyclic-GMP-AMP (3’3’-cGAMP), and 3’3’-cyclic-di-AMP (3’3’-CDA) were purchased from Invivogen and reconstituted in endotoxin-free water. Rabbit polyclonal antibodies against, IRF3 (1:1000), pIRF3 (S396, 1:1000), STING (1:1000), pSTING (S366, 1:100) pSTING (pS365, 1:1000), and Flag (1:1000) were purchased from Cell Signaling Technology. Mouse monoclonal anti-a-tubulin (1:4000) was purchased from Sigma-Aldrich.

#### Mouse Models

Mice were maintained at Stanford University in compliance with the Stanford University Institutional Animal Care and Use Committee (IACUC) regulations. All procedures were approved by the Stanford University Administrative Panel on Laboratory Animal Care (APLAC).

#### Flow Cytometry Analysis of Tumors

Tumor analysis was done as previously described^9^. The 7−9-week-old female BALB/c mice (Jackson Laboratories) were inoculated with 5 × 10^4^ 4T1-luciferase cells suspended in 50 μL of PBS. The cells were injected into the right fifth mammary fat pad. When tumor volume reached 100 ± 20 mm3, tumors were irradiated with 12 Gy using a 225 kVp cabinet X-ray irradiator with a 0.5 mm Cu filter (IC-250, Kimtron Inc.). Mice were anesthetized with a mixture of 80 mg/kg ketamine (VetaKet) and 5 mg/kg xylazine (AnaSed) prior to irradiation and were shielded with a 3.2 mm lead shield with 15 mm × 20 mm apertures to expose the tumors. Mice were then intratumorally injected with 100 μL of 100 μM neutralizing STING or nonbinding STING 24 h after irradiation. Mice were euthanized 24 h later, and the tumors were extracted. Following tumor extraction, the tumors were incubated at 37°C for 30 min in 10 mL of RPMI supplemented with 10% heat-inactivated FBS and 1% penicillin-streptomycin as well as 20 μg/mL DNase I type IV (Millipore) and 1 mg/mL collagenase from *Clostridium histolyticum* (Sigma-Aldrich). The samples were then passed through a 100 μm cell strainer (Sigma-Aldrich) to form a single-cell suspension. Red blood cells were lysed in 155 mM NH_4_Cl, 12 mM NaHCO_3_, and 0.1 mM EDTA for 5 min at room temperature. Samples were stained with LIVE/DEAD Fixable Blue Dead Cell Stain (Invitrogen) for 30 min. Samples were then fixed and permeabilized with either eBioscience Foxp3/Transcription Factor Staining Buffer Set (Invitrogen) or Fixation/Permeabilization Solution Kit (BD Biosciences). Samples were Fc blocked for 10 min using TruStain fcX (BioLegend) and then stained for 1 h. All samples were run on an Aurora analyzer (Cytek).

#### Isolation of Primary T cells

For human CD3^+^ T cells, buffy coat (Stanford Blood Center) was diluted 1:3 with PBS with 2% heat-inactivated FBS. Human T cells were then isolated from a diluted buffy coat using RosetteSep Human T Cell Enrichment Cocktail (StemCell Technologies) and SepMate tubes (StemCell Technologies) per manufacturer’s protocol. Isolated cells were cryopreserved in liquid nitrogen until further use. Purity of isolated cells was determined by flow cytometry to be >90%.

For mouse CD4+ and CD8+ T cells, spleens from 8-12 weeks old C57BL/6J mice were isolated and mashed through 70 μM strainer. Cell suspension was then pelleted and treated with 1X RBC Lysis Buffer (Invitrogen) for 15 min. Following lysis, cells were washed with PBS, and mouse CD4+ and CD8+ T cells were isolated using MojoSort Mouse CD4 T Cell Isolation Kit (BioLegend) and MojoSort Mouse CD8 T Cell Isolation Kit (BioLegend), respectively, per manufacturer’s protocol. Isolated cells were immediately activated for further use. Purity of isolated cells was determined by flow cytometry to be >90%.

#### Activation of Jurkat and primary T cells

Jurkat cells (1-2 × 10^6^ cells/mL) were activated with 5 ng/mL Phorbol 12-myristate 13-acetate (Cayman Chemical) and 500 ng/mL Ionomycin (Cayman Chemical) for 6 h. Primary human CD3^+^ T cells (1 × 10^6^ cells/mL) were activated with Dynabeads Human T-Activator CD3/CD28 (ThermoFisher) at 1:1 cell-to-bead ratio for 4 days. Primary mouse CD4+ and CD8+ T cells (1 × 10^6^ cells/mL) were activated with Dynabeads™ Mouse T-Activator CD3/CD28 (ThermoFisher) at 4:3 cell-to-bead ratio for 2 days.

#### CDN Killing Assay

For some experiments, U937 SLC19A1^-/-^ cell lines were pre-treated with 1 µg/mL of Doxycycline (Sigma-Aldrich) for 24 h to induce expression of SLC7A1 or SLC7A2 protein. U937 SLC19A1^-/-^ cell lines (5 × 10^5^ cells/ mL) and primary human CD3^+^ T cells (3 × 10^5^ cells/ mL) were treated with the indicated concentrations of CDN for 24 h, in the presence of 500 nM STF-1623^6^. To measure relative viability, cells were diluted 1:3 in PBS, and 10 µL of CellTiter-Glo (Promega) were added directly to 10 µL of diluted cells in a 384-well plate. The mixture was shaken gently and incubated at RT for 20 min. The luminescence was measured on a Tecan Spark with a 500 ms integration time. To measure relative count, cells were diluted 1:3 in PBS with 2 mM EDTA and 1:1000 DAPI (ThermoFisher). Cells were counted using Forward Scatter (FSC) and Side Scatter (SSC) gating, and the absence of DAPI staining on an AttuneNxt Flow Cytometer.

#### CDN Stimulation

For some experiments, U937 SLC19A1^-/-^ cell lines (1 × 10^6^ cells/mL), Jurkat cell lines (1 × 10^6^ cells/mL), and human and mouse primary T cells (1 × 10^6^ cells/mL) were pretreated with methotrexate (Sigma-Aldrich), folinic acid (Sigma-Aldrich), sulfasalazine (Sigma-Aldrich), DCPIB (Tocris Bioscience), for 10 min at the indicated concentration. For some experiments, U937 SLC19A1^-/-^ cell lines were pre-treated with 1 µg/mL of Doxycycline (Sigma-Aldrich) for 24 h to induce expression of SLC7A1 or SLC7A2 protein. Cells were treated with the indicated concentration of CDN for 2 h in a 5% CO_2_ incubator at 37°C, unless otherwise indicated. Following treatments, cells were collected, lysed with Laemmli Sample Buffer, and run on SDS-PAGE gels for Western blot analysis.

#### Jurkat CRISPR knockout library generation

Jurkat CRISPR knockout library line was created as previously described^37^. Briefly, a whole-genome library of exon-targeting sgRNAs were designed, with the goal of minimizing off-target effects and maximizing gene disruption. The top 10 sgRNA sequences for each gene were included in the library, along with thousands of safe-targeting and non-targeting negative controls. The library was cloned into a lentiviral vector, pMCB320, which also expresses mCherry and a puromycin resistance cassette. Jurkat cell line was transduced with a BFP-tagged Cas9 plasmid, selected clonally and Cas9 function was confirmed by the efficiency of knocking out GFP expression using a sgRNA sequence against GFP confirmed by using a sgRNA sequence against GFP. Jurkat-Cas9 cells were infected with the lentiviral library, and then selected with puromycin.

##### Live/Dead CRISPR screen

The Jurkat CRISPR knockout library line was activated with 5 ng/mL Phorbol 12-myristate 13-acetate (Cayman Chemical) and 500 ng/mL Ionomycin (Cayman Chemical) for 6 h. Cells were then grown in 4 spinner flasks (1 L), with 2 flasks serving as untreated controls and 2 flasks receiving cGAMP treatment. Throughout the screen all the samples were split daily to keep the cell density at 250 million cells per 200 mL, which corresponded to 1,000 cells per guide in the untreated samples. The experimental samples were treated with 2’3’-cGAMP starting at day 2 with enough cGAMP to reduce cell viability by 30% each day as compared to the control samples for 10 days. The initial treatment was 100 µM and was increased steadily to 300 µM by the final treatment day. At the end of the screen, the genomic DNA was extracted using a QIAGEN Blood Maxi Kit. The library was sequenced using a NextSeq 500/550 Mid Output v2 kit (Illumina). One experimental sample was dropped out of analysis due to failed sequencing QC. One experimental and two control samples were compared using casTLE^37^, available at https://bitbucket.org/ dmorgens/castle. The algorithm determines the likely effect size for each gene, as well as the statistical significance of this effect.

#### Generation of SLC7A1 Knock-Out Jurkat Line

LentiCRISPR v2 (Addgene) was used as the 3rd-generation lentiviral backbone for all knockout lines. A guide sequence targeting SLC7A1 (5’-GGGTGCTGGTGTCTACGTCC-3’) was cloned into the lentiviral backbone using the Lentiviral CRISPR Toolbox protocol from the Zhang Lab at MIT^39,40^. Lentiviral packaging plasmids (pHDM-G, pHDM-Hgmp2, pHDM-tat1b, and pRC/CMV-rev1b) were purchased from Harvard Medical School. To generate lentivirus, 500 ng of the lentiviral backbone plasmid containing the guide sequence and 500 ng of each of the packaging plasmids were transfected into HEK293T cells using FuGENE 6 transfection reagent (Promega).

Cell supernatant containing lentivirus was harvested after 48 h, and passed through a 0.45 μm filter. 1 mL of filtered supernatant was supplemented with 8 mg/mL Polybrene (Sigma-Aldrich) and added to 1 × 10^6^ Jurkat cells in a 12-well plate. Cells were spun at 700 xg for 30 min, and left to incubate overnight at 37°C. The next day, the virus containing media was removed and cells were resuspended in fresh media. Cells were selected 48 h later with 1 μg/ml Puromycin (Sigma-Aldrich) alongside control cells (uninfected) until all control cells died. Heterozygous knock-out population was assayed for CDN stimulation within 1 week after antibiotic selection.

#### CRISPR/Cas9 KO of Primary T Cells

For human T cells, negative (non-targeting) control and SLC7A1 (5’-GGGTGCTGGTGTCTACGTCC-3’) sgRNAs were purchased from Synthego and resuspended to 100 μM in TE buffer. Cas9 RNPs were formed by adding 1 μL of 61μM Alt-R S.p. Cas9 Nuclease V3 (IDT) to 1 μL of 100 μM sgRNA and incubating for 30 min at room temperature. Freshly activated CD3^+^ T cells were washed once with PBS, then resuspended in P3 Primary Cell nucleofector solution (Lonza) to a density of 1.5 × 10^6^ cells/ 20 μL. A volume of 20 μL of resuspended T cells was then added to the Cas9 RNPs, transferred to a Nucleocuvette stripwell, and nucleofected using program EO-115 on a 4-D Nucleofector X unit (Lonza).

Immediately after nucleofection, 150 μL of warm media was added to the cells. Cells were then then transferred to a 24-well plate containing 1 mL of RPMI, 10% heat-inactivated FBS, 1% penicillin-streptomycin and 300 U/mL rIL-2. At 24 h after nucleofection, cells were pelleted and resuspended in 2 mL of fresh media. At 72 h after transfection, cells were used for CDN stimulation assays and genomic DNA was isolated to measure the knockout efficiency. The knockout efficiency was determined by amplifying the region of genomic DNA surrounding sgRNA target sites, performing Sanger sequencing, and using the sequencing trace to estimate knockout efficiency through ICE (Synthego) analysis.

For mouse T cells, Rosa26 and SLC7A1 (5’-ATTTTCACGGGCCACGGCAC-3’) sgRNAs were purchased from Synthego and resuspended to 100 μM in TE buffer. Cas9 RNPs were formed by adding 0.6 μL of 61μM Alt-R S.p. Cas9 Nuclease V3 (IDT) to 1 μL of 100 μM sgRNA and incubating for 30 min at room temperature. Freshly activated CD4+ and CD8+ T cells were washed once with PBS, then resuspended in P3 Primary Cell nucleofector solution (Lonza) to a density of 1.5 × 10^6^ cells/ 20 μL. A volume of 20 μL of resuspended T cells was then added to the Cas9 RNPs, transferred to a Nucleocuvette stripwell, and nucleofected using program DN-100 on a 4-D Nucleofector X unit (Lonza). Immediately after nucleofection, 150 μL of warm media was added to the cells. Cells were then then transferred to a 24-well plate containing 1 mL of RPMI, 10% heat-inactivated FBS, 1% penicillin-streptomycin, 50 µM 2-Mercaptoethanol, 2 mM L-Citrulline (TCI Chemicals), and 300 U/mL rIL-2. At 24 h after nucleofection, cells were pelleted and resuspended in 2 mL of fresh media. At 72 h after transfection, cells were used for CDN stimulation assays and genomic DNA was isolated to measure the knockout efficiency. The knockout efficiency was determined by amplifying the region of genomic DNA surrounding sgRNA target sites, performing Sanger sequencing, and using the sequencing trace to estimate knockout efficiency through ICE (Synthego) analysis.

#### Electroporation of 2’3’-cGAMP

U937 SLC19A1^-/-^ cell lines were pelleted and resuspended in nucleofector solution (90 mM Na_2_HPO_4_,90 mM NaH_2_PO_4_, 5 mM KCl, 10 mM MgCl_2_, 10 mM sodium succinate) with 100 nM 2’3’-cGAMP to a density of 1 × 10^6^ cells/mL. A volume of 100 μL of cells was then transferred to a 0.2 cm electroporation cuvette and electroporated with program U-013 on a Nucleofector IIb device. Immediately after nucleofection, 500 μL of warm media was added to the cells. Cells were then transferred to a 24-well plate containing an additional 500 μL of media and incubated at 37°C for 2 h. Following this, cells were collected, lysed with Laemmli Sample Buffer, and run on SDS-PAGE gels for Western blot analysis.

Mouse primary CD4+ and CD8+ T cells were pelleted and resuspended in P3 Primary Cell Nucleofector Solution (Lonza) with 100 nM 2’3’-cGAMP to a density of 1 × 10^6^ cells/mL. A volume of 20 μL of cells was then transferred to the Nucleocuvette stripwell and electroporated with program DN-100 on a 4-D Nucleofector X unit (Lonza). Immediately after nucleofection, 150 μL of media was added to the cells. Cells were then transferred to a 24-well plate containing an additional 850 μL of media and incubated at 37°C for 3 h. Following this, cells were collected, lysed with Laemmli Sample Buffer, and run on SDS-PAGE gels for Western blot analysis.

#### SLC7A1 Protein Purification

To express SLC7A1 protein, Expi293 cells were cultured at a density of 3 × 10^6^ cells/mL. Cells were then transfected with pcDNA3.1-FLAG-HRV3C-SLC7A1 plasmid was transfected with FectoPro transfection reagent (VWR) and supplemented with 4 g/L of D-glucose (Sigma Aldrich) and 3 mM of valproic acid (Sigma-Aldrich).

To purify SLC7A1 protein, Expi293 cells were harvested at 72 h post-transfection, snap-frozen in liquid nitrogen, and thawed on ice in 20 mL of hypotonic buffer (10 mM HEPES pH 7.5, 25 mM NaCl). Cell mixture was then mechanically dounced, and 1 eq. volume of 2x isotonic buffer (50 mM HEPES pH 7.5, 600 mM NaCl) and EDTA-free cOmplete protease inhibitor cocktail (Roche) were added. Mixture was spun at 100,000 xg for 40 min, and the resulting pellet (membrane fraction) was solubilized for 1 h at 4°C in solubilization buffer (25 mM HEPES pH 7.5, 300 mM NaCl, 1% DDM/0.1% CHS). Protein mixture was then spun at 100,000 xg for 40 min, and the resulting supernatant containing membrane proteins was collected. Anti-DYKDDDDK affinity resin (ThermoFisher) were incubated overnight with the membrane proteins, washed with washing buffer (25 mM HEPES pH 7.5, 500 mM NaCl, 2 mM CaCl_2_, 0.1% DDM/0.01% CHS), and SLC7A1 proteins were eluted from the resin with 200 μg/mL FLAG Peptide (MedChemExpress) in elution buffer (25 mM HEPES pH 7.5, 150 mM NaCl, 5 mM EDTA, 0.1% DDM/0.01% CHS). Eluted SLC7A1 proteins were dialyzed into PBS with 0.04% DDM/ 0.004% CHS. Samples from every purification steps were run on SDS-PAGE gel and stained with Coomasie total protein stain or blotted for anti-Flag on Western Blot.

#### Synthesis of DncV

Rosetta cells expressing pTB146-His-SUMO-DncV were grown in 2xYT medium with 100 mg/mL ampicillin and induced with 0.5 mM IPTG when OD_600_ reached 1, and then were grown overnight at 16°C. All subsequent procedures using proteins and cell lysates were performed at 4°C. Cells were pelleted and lysed in 20 mM HEPES pH 7.5, 400 mM NaCl, 10% glycerol, 10 mM imidazole, 1 mM DTT, and EDTA-free, cOmplete protease inhibitor cocktail (Roche). Cell lysate was then cleared by ultracentrifugation at 50,000 x g for 1 h. The cleared supernatant was incubated with HisPur cobalt resin (ThermoFisher Scientific; 1 mL resin per 1 L bacterial culture) for 1 hour. Cobalt resin was then washed with 20 mM HEPES pH 7.5, 1 M NaCl, 10% glycerol, 10 mM imidazole, and 1 mM DTT. Protein was eluted from resin with 300 mM imidazole in 20 mM HEPES pH 7.5, 400 mM NaCl, and 1 mM DTT. Fractions containing His-SUMO-DncV were pooled, concentrated, and dialyzed against 20 mM HEPES pH 7.5, 400 mM NaCl, 1 mM DTT, and then snap-frozen in aliquots for future use.

#### Synthesis of 3’3’-cGAMP diazirine probe

To synthesize 3’3’-cGAMP diazirine probe, 1 mM of 6’Cl ATP (Jena Biosciences) and 1 mM of GTP (Sigma-Aldrich) were mixed in 20 mM of Tris-HCl at pH 7.4 and 20 mM of MgCl_2_. Subsequently, 200 nM of purified DncV was added, and the reaction mixture was stirred vigorously at 37°C for 1h. The reaction’s progress was monitored via Thin Layer Chromatography (TLC) and LC-MS (Waters). The product 6’-Cl-2’3’-cG(A)MP was purified using HPLC on a PFP Poroshell 120 column (Agilent Technologies, Inc.) with Acetonitrile/0.1% Formic Acid as the mobile phase. Purified product was subsequently lyophilized and resuspended in water. 6’-Cl-2’3’-cG(A)MP was then mixed with 5 eq. of amino diazirine alkyne and 20 eq. of triethylamine, and the reaction was stirred at 40°C overnight. The product was purified with HPLC as previously described in 2’3’-cGAMP synthesis step^29^.

#### Diazirine crosslinking competition assay

cGAMP diazirine probe (500 μM) and 2’3’- and 3’3’-cGAMP (500 μM for low concentration, 2 mM for high concentration; water for controls) were mixed in binding buffer (20 mM Tris-HCl pH 7.4, 200 mM NaCl, 1 mM MgCl_2_). 0.1 mg/mL of purified human SLC7A1 protein was subsequently added to a volume of 10 μL, and the mixture was incubated for 10 min at RT. The mixture was then crosslinked at 365 nm with UVP crosslinker CL-3000 (Analytik Jena US) on ice for 20 minutes. To a crosslinked reaction mixture, 0.1 mM of R110 azide (MedChemExpress) was added into a pre-mixed complex of 1 mM of CuSO_4_ and 5 mM BTTAA (Sigma-Aldrich), which were mixed with 5 mM of sodium ascorbate to a total volume of 15 μL. The click reaction mixture was incubated for 1-2 h at RT. The reaction was quenched with the addition of 15 μL 2x reducing Laemmli Sample Buffer. Sample was then assayed by Western blot for R110 fluorescence and Flag signal.

#### Generation of Doxycycline Inducible Cell Lines

Lentiviral packaging plasmids (pHDM-G, pHDM-Hgmp2, pHDM-tat1b, and RC/CMV-rev1b) were purchased from Harvard Medical School. To generate lentivirus, 500 ng of lentiviral plasmid encoding doxycycline inducible transporters and 500 ng of each of the packaging plasmids were transfected into HEK 293T cells with FuGENE 6 transfection reagent (Promega). Cell supernatant containing lentivirus was harvested after 48 h, and passed through a 0.45 μm filter. To create the inducible cell lines in U937 SLC19A1^−/−^,^29^1 mL of filtered supernatant was supplemented with 8 mg/mL Polybrene (Sigma-Aldrich) and added to 5 × 10^5^ cells in a 12-well plate. Cells were spun at 700 xg for 30 min, and left to incubate overnight at 37°C. The next day, the virus containing media was removed and cells were resuspended in fresh media. Cells were selected 48 h later with 5 μg/ml Blasticidin (Sigma-Aldrich) alongside control cells (uninfected) until all control cells died.

#### [^3^H]-Arginine Uptake Assay

U937 SLC19A1^-/-^ cell lines (1 × 10^7^ cells) were treated with 21 nM [^3^H]-L-Arginine (Perkin Elmer) in RPMI SILAC buffer (ThermoFisher) for 1 min. For some experiments, L-Lysine (Sigma-Aldrich) or 2’3’-cGAMP was added at the same time as [^3^H]-L-Arginine at the indicated concentrations. Cold PBS was then added to cells to halt import. Following two more washes in cold PBS, cells were lysed in 500 μL of 400 mM NaOH and heated at 65°C for 45 min to completely dissolve the lysate. Then, 400 μL of cell lysate was used to count ^3^H signal on a scintillation counter and 25 μL were used in a bicinchoninic acid assay (Thermo Scientific) to quantify protein levels for normalization.

#### Bulk RNA-Seq

U937 SLC19A1^-/-^ cell lines (5 × 10^6^ cells) were treated with 1 µg/mL of Doxycycline (Sigma-Aldrich) for 24 h to induce expression of SLC7A1 or SLC7A2 protein. Cells were then harvested and lysed in TRIzol reagent (ThermoFisher). Total RNA was then purified using Direct-zol RNA purification kit (Zymo Research). RNA from different samples (n = 2 biological replicates) were then shipped to sequencing service provider Innomics for QC testing and sequencing on DNBSEQ platform. FASTQs were demultiplexed, mapped onto human reference genome (GRCh38.p13), and differential gene expression were generated using the DESeq2 method^36^ by sequencing service provider Innomics.

#### Modeling the cGAMP and Arginine Binding Sites in SLC7A1/2

AlphaFold structure of human SLC7A1 (UniProt ID P30825) was minimized using Schrödinger Maestro using standard protein preparation parameters at all default settings at pH 7.4. Binding site detection was run to identify possible pocket sites on SLC7A1 and the predicted pocket scoring highest and occurring on the extracellular-facing region of the structure was selected for receptor grid preparation (all standard parameters). All possible conformations of 2’,3’-cGAMP at pH 7.4 were prepared using LigPrep, and the set of structures docked against the SLC7A1 receptor grid using Glide at standard precision.

The minimized SLC7A1 structure was aligned to the crystal structure of arginine-bound *Geobacillus* LAT (PDB: 6F34) and the ligand was used to create a composite SLC7A1–arginine structure specifying a candidate amino acid binding and transport site. The site was similarly used for receptor grid preparation, and re-docked with prepared arginine conformations for validation of the model, before cGAMP structures were separately docked.

#### cGAMP Uptake Experiment

U937 SLC19A1^-/-^ tet-SLC7A2-Flag cells (1 × 10^7^ cells) were treated with 1 µg/mL of Doxycycline (Sigma-Aldrich) for 24 h. Cells were then treated with 10 µM cGAMP at 37°C for 15 min in serum-free RPMI SILAC with 500 nM of STF-1623. Cold PBS was then added to cells to halt import. Following 3 more washes, cells were lysed in T-PER protein extraction buffer (ThermoFisher), and heated at 80°C for 10 min to release STING-bound cGAMP. Intracellular cGAMP level was measured using 2’3’-cGAMP ELISA kit (Cayman Chemical) and normalized to protein level from the resulting supernatant.

#### Quantification and Statistical Analysis

All statistical analyses were performed using GraphPad Prism 10. All p values were calculated using an unpaired t test with Welch’s correction. For all experiments involving Western blots, densitometric measurements of protein bands were made using ImageJ 2.30.

---

**Supplementary Figure 1.**
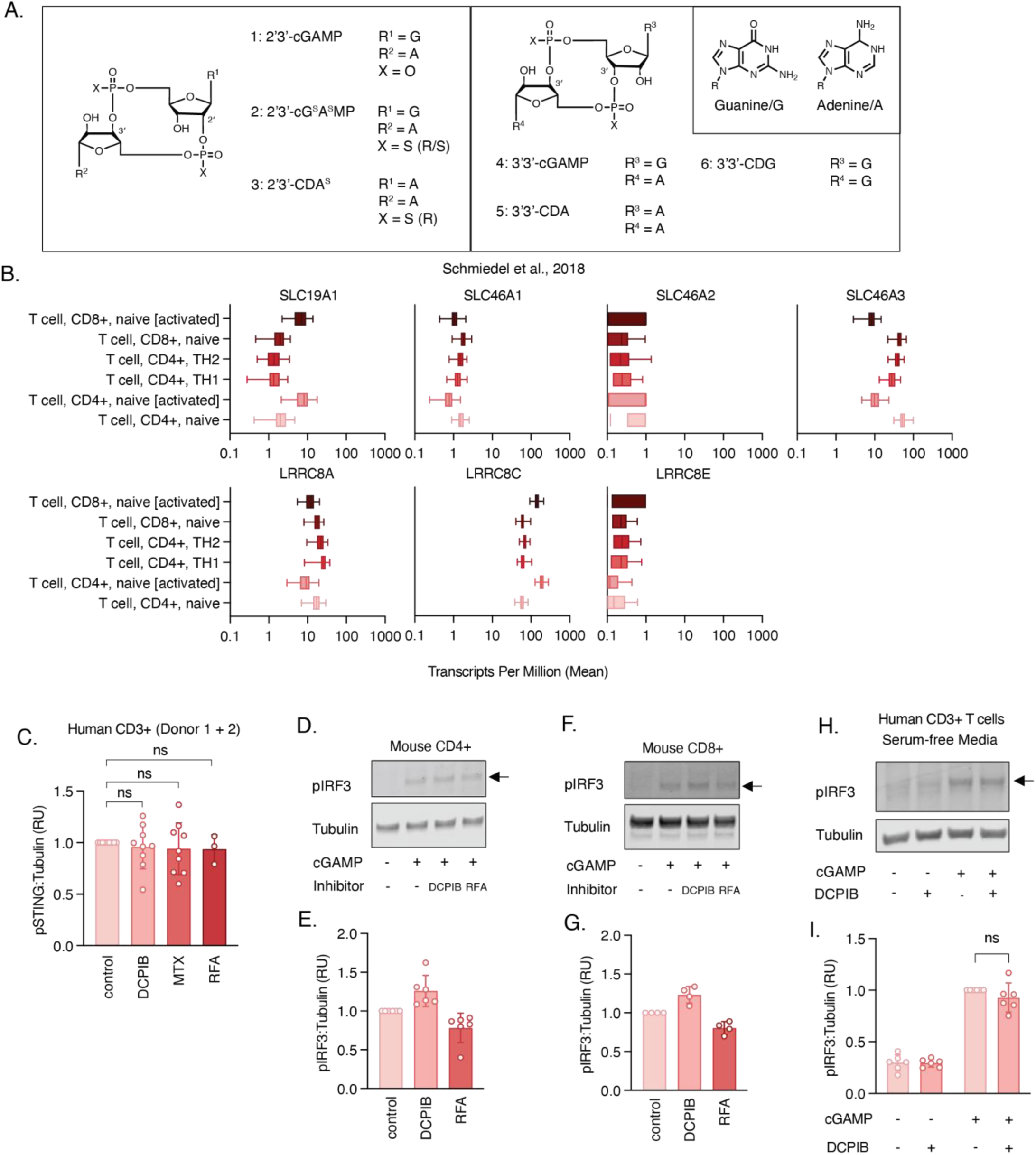
**Related to Figure 1.** (A) Structures of mammalian (1), synthetic (2-3) and bacterial (4-6) cyclic dinucleotides. (B) Human transcript database adopted from Schmiedel et al., 2018 for known cGAMP importers in different human T cell subsets. (C) Combined quantification of Western blots from Figure 1E. Significance are calculated by unpaired, two-tailed t-test. Data are shown as the mean +/- SD. (D and E) Activated primary mouse CD4+ T cells were treated with 50 µM cGAMP for 3 h, and with or without 20 µM DCPIB or 0.5 mM reduced folic acid (RFA), and and signaling was assessed by Western blot (n = 6 biological replicates). One representative Western blot is shown in (E) and quantification of multiple blots is shown in (F). Data are shown as the mean +/- SD. (F and G) Activated primary mouse CD8+ T cells were treated with 50 µM cGAMP for 3 h, and with or without 20 µM DCPIB or 0.5 mM reduced folic acid (RFA), and and signaling was assessed by Western blot (n = 4 biological replicates). One representative Western blot is shown in (G) and quantification of multiple blots is shown in (H). Data are shown as the mean +/- SD. (H and I) Activated human primary T cells from two healthy donors were treated with 50 µM cGAMP for 2 h, and with or without 20 µM DCPIB, in serum-free media, and and signaling was assessed by Western blot (n = 6 biological replicates). One representative Western blot from donor 2 is shown in (H) and quantification of multiple blots are shown in (I). Data are shown as the mean +/- SD. Significance are calculated by unpaired, two-tailed t-test.

**Supplementary Figure 2.**
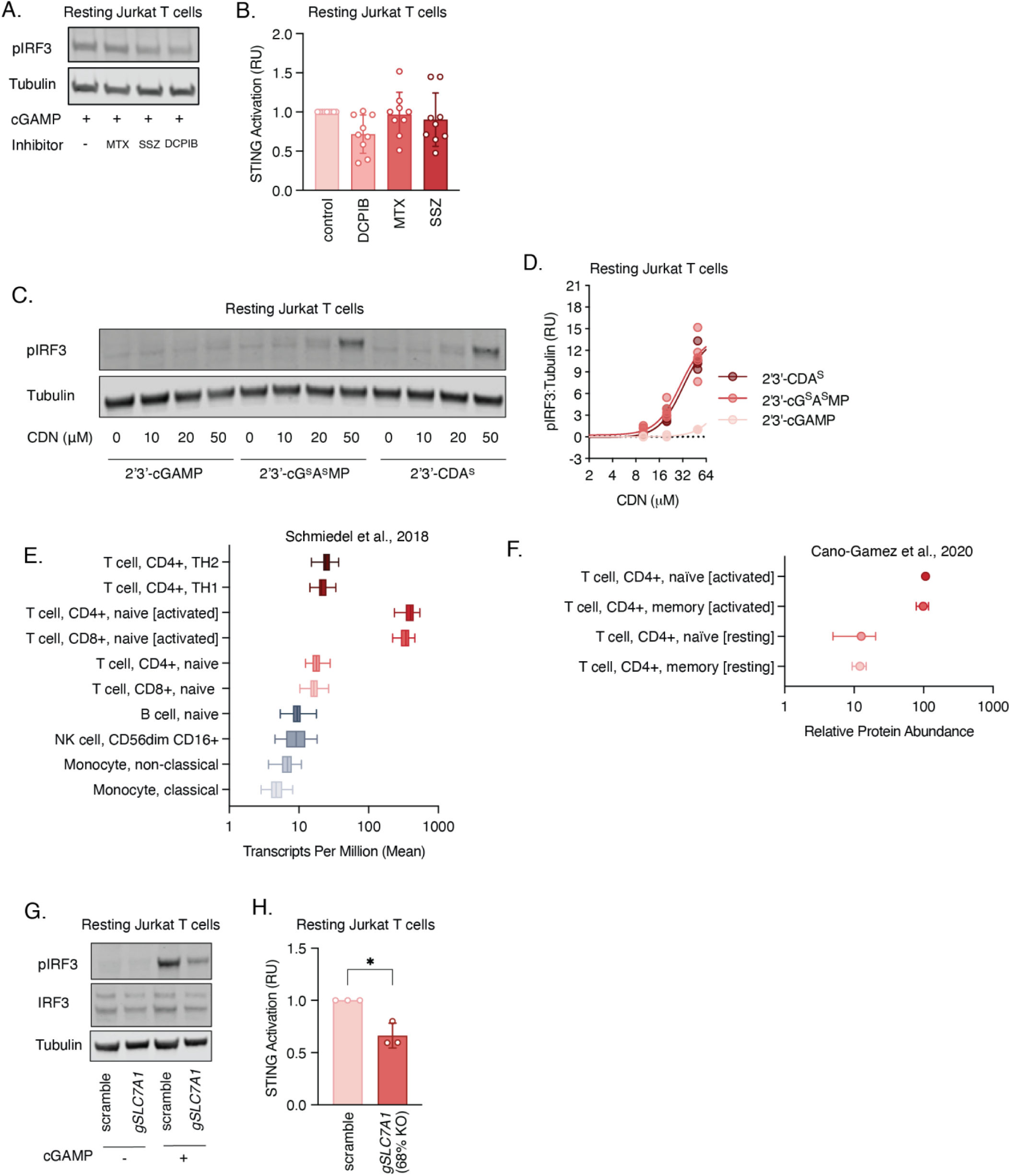
**Related to Figure 2.** (A and B) Jurkat cells were treated with 50 µM cGAMP for 2 h, and with or without 0.5 mM methotrexate (MTX), 1 mM sulfasalazine (SSZ) or 20 µM DCPIB, and STING activation (pIRF3:Tubulin or pSTING:Tubulin) was assessed by Western blot (n = 9 biological replicates). One representative Western blot is shown in (A) and quantification of multiple blots is shown in (B). Data are shown as the mean +/- SD. (C and D) Jurkat cells were treated with increasing concentration of 2’3’-cGAMP, 2’3’-cG^S^A^S^MP or 2’3’-CDA^S^ for 2 h, and signaling was assessed by Western blot of pIRF3 or pSTING (n = 3 - 5 biological replicates as indicated). One representative Western blot is shown in (C) and quantification of multiple blots is shown in (D). Data are shown as the mean +/- SD. (E) Human transcript database adopted from Schmiedel et al., 2018 for SLC7A1 expression in different immune cell subsets. (F) Human proteomic database adopted from Cano-Gamez et al., 2020 for SLC7A1 expression in resting and activated T cell subsets. (G and H) Jurkat cells were transduced with Cas9 along with non-targeting (scramble) or SLC7A1-targeting sgRNA, were treated with 75 uM cGAMP for 2h, and STING activation (pIRF3:Tubulin or pSTING:Tubulin) was assessed by Western blot (n = 3 biological replicates). Knock-out score was determined by ICE (Synthego) to be 68%. One representative Western blot is shown in (G) and quantification of multiple blots is shown in (H). Significance calculated by unpaired, two-tailed t-test. Data are shown as mean +/- SD. *p < 0.05

**Supplementary Figure 3.**
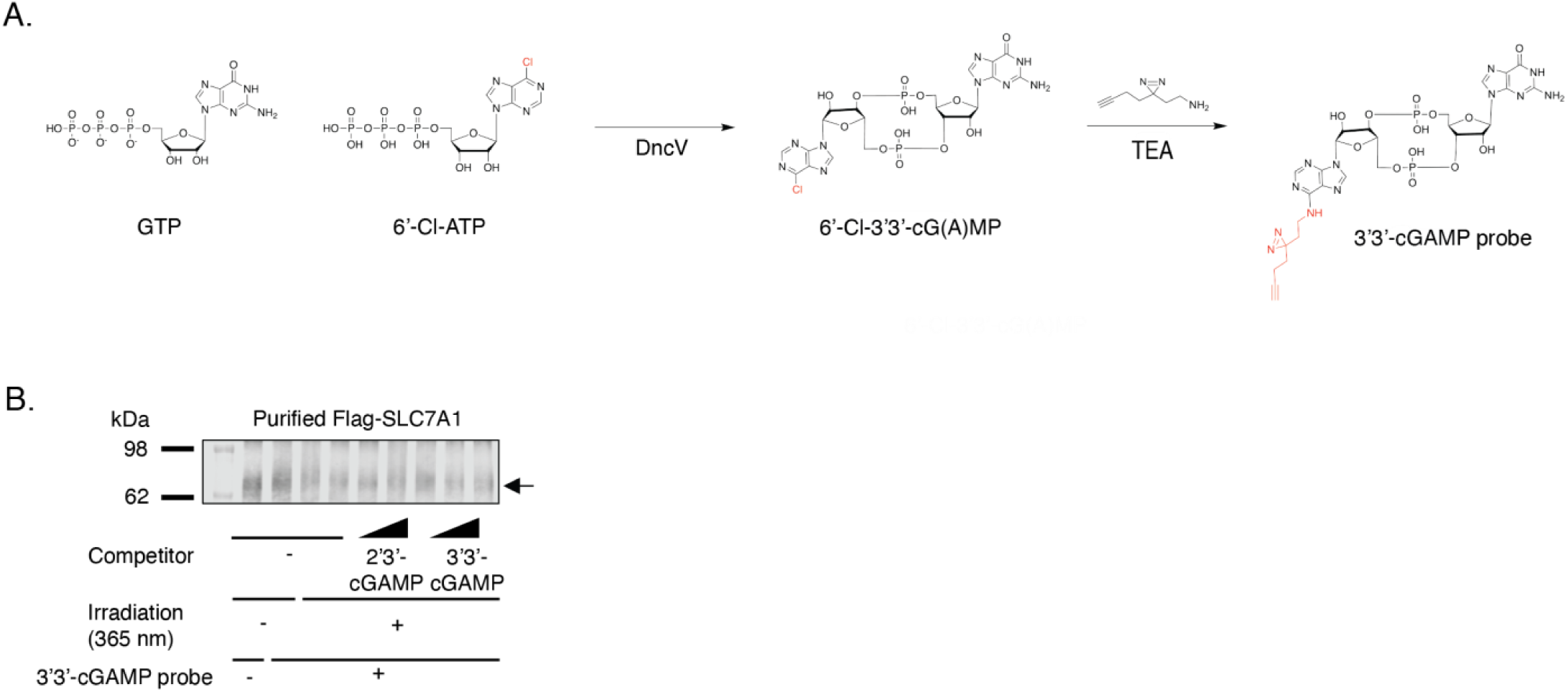
**Related to Figure 3.** (A) Synthesis schematic of 2’3’- and 3’3’ cGAMP-Diazirine probe. (B) Flag blot of purified FLAG-SLC7A1 from (Figure 3H).

**Supplementary Figure 4.**
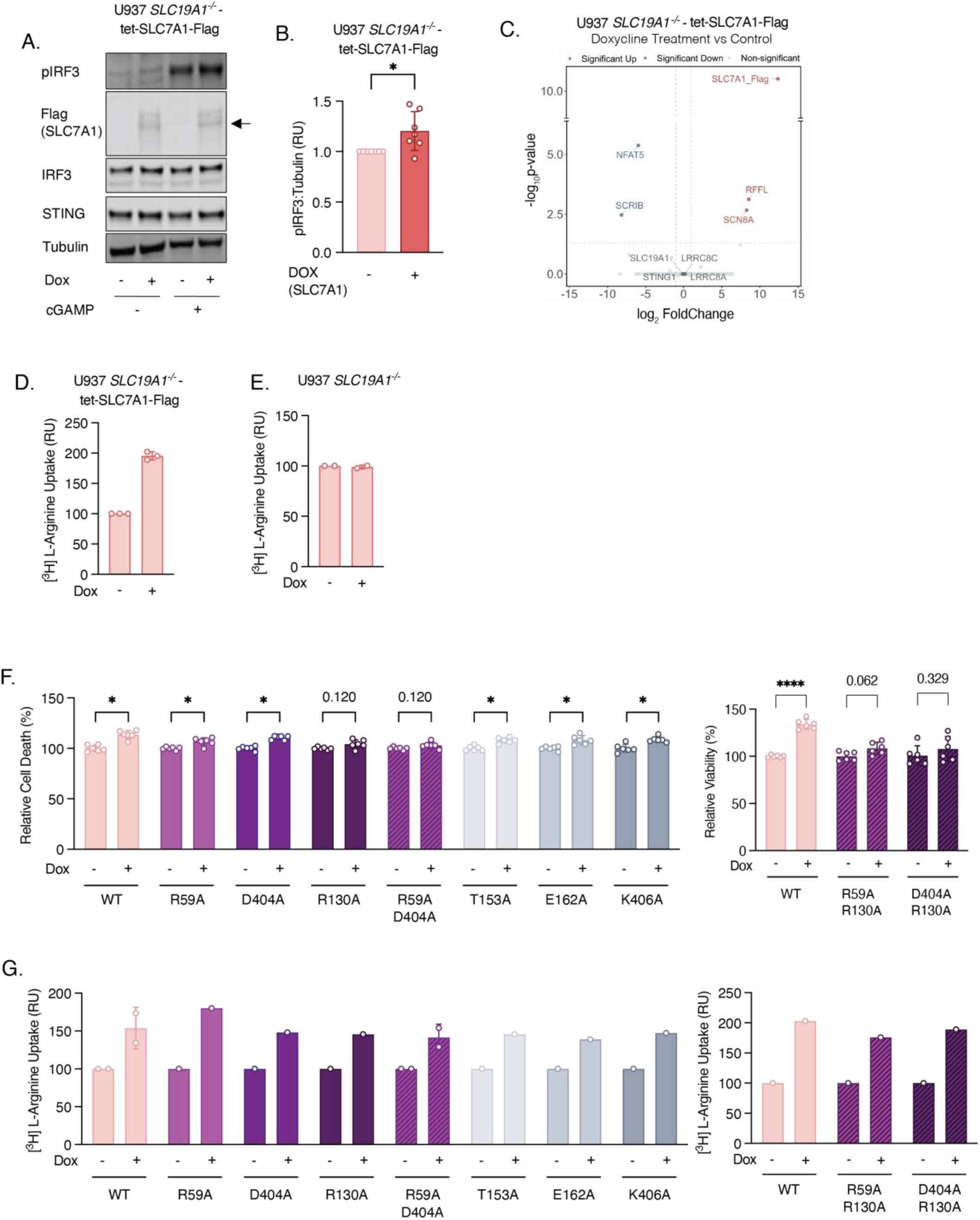
**Related to Figure 4.** (A and B) U937 SLC19A1^-/-^ -tet-SLC7A1-Flag cells were incubated with or without 1 µg/mL doxycycline (dox) for 24 h. Cells were then treated with 75 µM cGAMP for 2 h, and signaling was assessed by Western blot (n = 7 biological replicates). One representative Western blot is shown in (A) and quantification of multiple blots is shown in (B). Significance calculated by unpaired, two-tailed t-test. Data are shown as mean +/- SD. *p < 0.05 (A) Differential gene analysis (DESeq2) of U937 SLC19A1^-/-^ -tet-SLC7A1-Flag cells incubated with or without 1 µg/mL doxycycline (dox) for 24 h (n = 2 biological duplicate). Significantly enriched or depleted coding mRNAs and known extracellular cGAMP/STING signaling elements are annotated. (B) U937 SLC19A1^-/-^ -tet-SLC7A1-Flag cells were incubated with or without 1 µg/mL doxycycline (dox) for 24 h. Cells were then treated with 21 nM ^3^H-Arginine for 1 min (n = 3 biological replicates), and radioactivity was measured using a scintillation counter and normalized to protein amount. Data are shown as the mean +/- SD. (C) U937 SLC19A1^-/-^ cells were incubated with or without 1 µg/mL doxycycline (dox) for 24 h. Cells were then treated with 21 nM ^3^H-Arginine for 1 min (n = 2 biological replicates), and radioactivity was measured using a scintillation counter and normalized to protein amount. Data are shown as the mean +/- SD. (D) U937 SLC19A1^-/-^ -tet-SLC7A1 (WT, single or double mutant)-Flag cells were incubated with or without 1 µg/mL doxycycline (dox) for 24 h. Cells were then treated with 20 µM 2’3’-cGAMP for 24 h in the presence of ENPP1 inhibitor, STF-1623. Relative viability was measured using Cell Titer Glo Assay (n = 6 biological replicates, as indicated). Significance calculated by unpaired, two-tailed t-test. Data are shown as the mean +/- SD. *p<0.05. (E) U937 SLC19A1^-/-^ -tet-SLC7A1 (WT, single or double mutant)-Flag cells were incubated with or without 1 µg/mL doxycycline (dox) for 24 h. Cells were then treated with 21 nM ^3^H-Arginine for 1 min (n = 1-2 biological replicates, as indicated), and radioactivity was measured using a scintillation counter and normalized to protein amount.

**Supplementary Figure 5.**
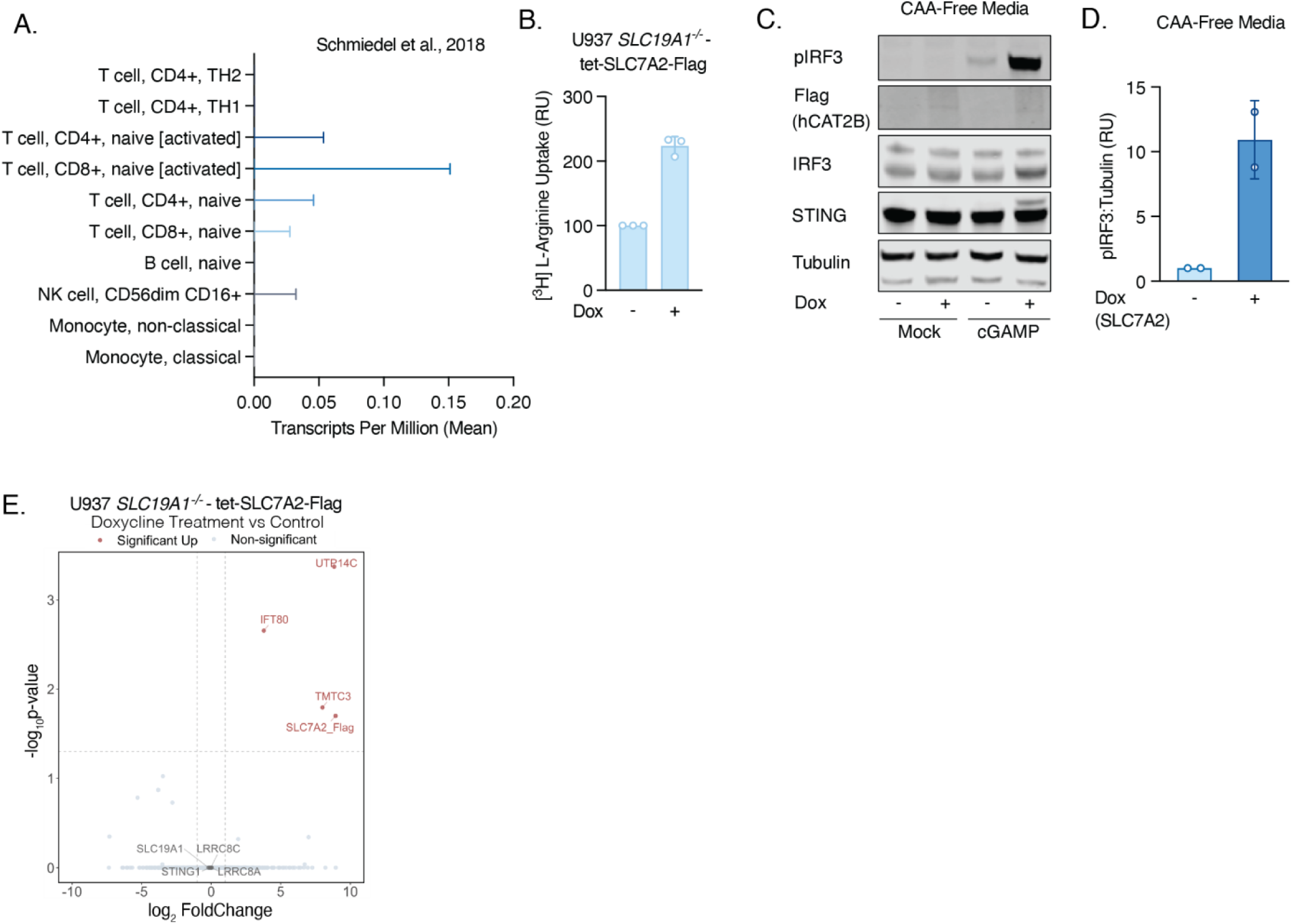
**Related to Figure 5.** (A) Human transcript database adopted from Schmiedel et al., 2018 for SLC7A2 expression in different immune cell subsets. (B) U937 SLC19A1^-/-^ -tet-SLC7A2-Flag cells were incubated with or without 1 µg/mL doxycycline (dox) for 24 h. Cells were then treated with 21 nM ^3^H-Arginine for 1 min (n = 3 biological replicates), and radioactivity was measured using a scintillation counter and normalized to protein amount. Data are shown as the mean +/- SD. (C and D) U937 SLC19A1^-/-^ -tet-SLC7A2-Flag cells were incubated in the absence of all cationic amino acids, and with or without 1 µg/mL doxycycline (dox) for 24 h. Cells were then treated with 75 µM cGAMP for 2 h, and signaling was assessed by Western blot (n = 2 biological replicates). One representative Western blot is shown in (C) and quantification of multiple blots is shown in (D). Data are shown as mean +/- SD. (E) Differential gene analysis (DESeq2) of U937 SLC19A1^-/-^ -tet-SLC7A2-Flag cells incubated with or without 1 µg/mL doxycycline (dox) for 24 h (n = 2 biological duplicate). Significantly enriched or depleted coding mRNAs and known extracellular cGAMP/STING signaling elements are annotated.

